# High-Speed AFM Reveals Ligand-Dependent Supramolecular Switching of Human Phosphofructokinase-1

**DOI:** 10.64898/2026.05.08.723867

**Authors:** Shuangyu Luo, Arnav Patil, Nicholas Primanis-Erickson, Xiaoding Jiang, Bradley A. Webb, Ku-Lung Hsu, Yi-Chih Lin

**Author notes:** Correspondence to: Yi-Chih Lin.

## Abstract

Phosphofructokinase-1 (PFK-1) catalyzes the ATP-dependent conversion of fructose-6-phosphate (F6P) to fructose-1,6-bisphosphate (F1,6BP), the first committed step of glycolysis. Beyond classical allostery, the liver isoform PFKL forms higher-order assemblies, but how ligand binding redirects intermolecular interactions remains unclear. Here we use high-speed atomic force microscopy (HS-AFM), topology-based AFM image simulations, and molecular dynamics (MD) simulations to define ligand-dependent assembly switching of human PFKL. Wild-type PFKL (PFKL WT) forms lattice-like assemblies under APO and ATP conditions, whereas coordinated ATP and F6P loading redirects assembly toward filaments in a ligand-order-dependent manner. APO-PFKL WT also forms lattice-like assemblies on or near membrane-supported surfaces, suggesting that interfacial environments influence where assembly initiates. The filament-defective N702T mutant forms ordered double-layer lattices and preserves this geometry under APO, ATP, and ATP+F6P conditions. MD simulations suggest that lattice stabilization arises from distributed inter-tetramer contacts. These findings define a structural framework for ligand-dependent supramolecular regulation of glycolysis.

## Introduction

Glycolysis is a core metabolic pathway that must rapidly adjust flux in response to changing cellular demands.^1^ Phosphofructokinase-1 (PFK1) catalyzes the ATP-dependent phosphorylation of fructose-6-phosphate (F6P) to fructose-1,6-bisphosphate (F1,6BP)^1,2^, the first committed step of glycolysis and a major rate-controlling reaction in the pathway. In mammals, three PFK1 isoforms, including liver (PFKL), platelet (PFKP), and muscle (PFKM) have been identified. These isoforms are expressed at different levels across tissues; many tissues express more than one isoform, whereas some tissues are enriched for a predominant isoform, such as PFKM in skeletal muscle and PFKL in hepatocytes.^3^ Among them, PFKL is unique in that its functional tetramers can self-assemble into filamentous structures in vitro.^4-6^ Because the liver plays a central role in maintaining glucose homeostasis^7^, PFKL self-assembly may represent an additional regulatory layer superimposed on classical allostery, allowing glycolytic flux to respond rapidly to changing metabolic demand.^8-10^

Multiple metabolic enzymes have been shown to assemble into filamentous structures, which could regulate their functions in the associated biological pathways.^8,11-13^ In some systems, filamentation correlates with reduced enzymatic activity^14,15^, whereas in others it stabilizes active conformations^16^, indicating that assembly can modulate function in multiple ways. However, a major gap in the field is that metabolic enzymes in cells often appear as spatially heterogeneous assemblies, including filaments, puncta, and higher-order clusters, yet whether these structures represent a single structural continuum or distinct functional states with different regulatory consequences remains unclear. This question is particularly relevant for glycolytic enzymes that must rapidly adapt activity without transcriptional regulation. For PFKL, ligands are known to alter enzymatic conformation and activity, but it remains unclear how ligand binding modulates intermolecular interactions to redirect higher-order self-assembly.

Consistent with tissue-dependent isoform expression, PFK1 can form homo- or heterotetramers depending on cellular context, including PFKL/PFKM-containing heterotetramers in erythrocytes.^3,17^ In hepatocytes, where PFKL is the predominant isoform, PFKL is expected to primarily form homotetramers. Each monomeric unit of PFKL consists of a catalytic domain and a regulatory domain bridged by a flexible linker domain (**Fig. 1A-1B and Fig. S1**).^6,18^ The PFKL tetramer contains catalytic sites that support F6P phosphorylation and regulatory sites through which allosteric ligands shift the conformational equilibrium between inactive (T-) and active (R-) states.^19,20^ In previous studies, PFKL tetramers in the R-state were shown to self-assemble into filaments in vitro through an interface involving residue N702 between adjacent tetramers (**Fig. 1C**).^5,6^ The N702T mutation, designed by substituting PFKL Asn702 with the threonine found at the corresponding position in the platelet isoform PFKP, prevents filament formation and was therefore used to probe the structural basis of PFKL filament assembly. These findings established residue N702 as a key component of the filament-forming interface, but they did not determine whether PFKL higher-order assembly is limited to filaments or can access alternative supramolecular states.

**Figure 1.**
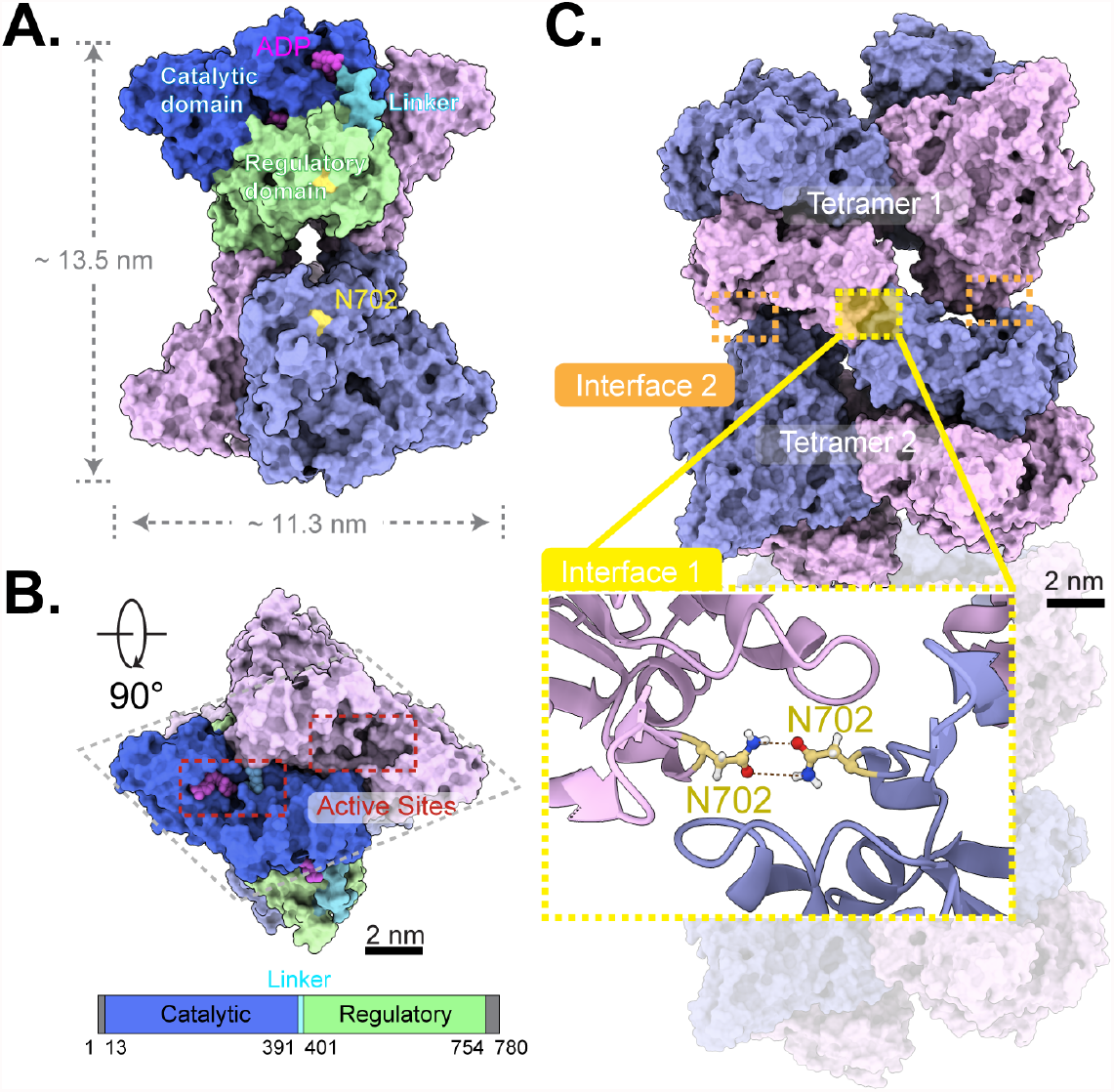
Structural organization of human PFKL and the filament-forming interface. **(A-B)** Structure of PFKL tetramer in its functionally active R-state (PDB: 8W2G) bound to F1,6BP, F6P, and ADP. One monomer is colored to indicate the catalytic domain (blue), interdomain linker (cyan), and regulatory domain (green). The PFKL tetramer has projected dimensions of approximately ∼13.5 nm × ∼11.3 nm when viewed from the side of the long axis. Catalytic and regulatory ligand-binding sites are distributed across the tetramer, allowing substrates and allosteric effectors to shift the conformational equilibrium between active (R-) and inactive (T-) states. **(C)** Filament architecture of R-state PFKL (PDB: 8W2I). Adjacent tetramers interact through an interface involving residue N702. Two additional tetramer-tetramer interfaces are indicated (orange boxes). This previously defined N702-containing interface provides the structural reference for testing whether ligand-dependent PFKL assembly is restricted to the previously described filament pathway or can access alternative intermolecular interfaces.

In cultured mammalian cells, PFKL forms punctate structures predominantly localized near the plasma membrane. Recent work further shows that PFKL WT localizes to lamellipodia in migrating breast cancer cells, whereas a filament-incompetent N702T mutant shows reduced lamellipodial recruitment and impaired directional sensing, supporting a functional role for higher-order PFKL organization in spatially localized cell behavior.^21^ PFKL has also been identified in a multienzyme glucose-metabolism complex in living human cells, supporting the view that glycolytic regulation can involve mesoscale spatial organization of enzymes rather than only regulation of isolated catalytic units.^22^ Recent development of a covalent PFKL activator that stabilizes the R-state tetramer and suppresses tumor growth further highlights PFKL as a structurally regulatable metabolic target.^23^ However, these studies do not define the structural relationship between puncta, lamellipodial PFKL assemblies, and filament formation, or whether distinct ligand conditions can bias PFKL toward different supramolecular architectures. Thus, a structural description connecting ligand state, intermolecular interactions, and assembly architecture is required to understand how enzyme organization contributes to metabolic regulation.

High-speed atomic force microscopy (HS-AFM)^24,25^ is a biophysical technique with high spatial and temporal resolution for studying biomolecular self-assembly in real time under near-physiological conditions, including the formation of annexin-V lattices on membranes^26^, septin filaments^27,28^, amyloid fibrils^29^, and ESCRT-III spirals^30^. To directly determine how ligand state modulates PFKL higher-order organization, we use HS-AFM to visualize PFKL assembly dynamics in real time. We first define the assembly behavior of wild-type PFKL (PFKL WT), showing that APO and ATP-bound conditions support lattice-like assemblies, whereas coordinated ATP and F6P binding redirects PFKL toward filament formation^5,6,31^. We then use the filament-defective N702T mutant as a mechanistic perturbation to test whether disruption of the previously described filament-forming interface abolishes higher-order assembly. Instead, PFKL N702T forms a highly ordered double-layer lattice, revealing that PFKL can access an alternative assembly state distinct from the filament pathway. HS-AFM resolves the surface topography of the PFKL N702T lattice, and comparison with topology-based AFM image simulations defines the molecular packing geometry of tetramers within the lattice. Molecular dynamics (MD) simulations further suggest that lattice stabilization arises from distributed inter-tetramer interactions rather than from the N702-dependent filament interface alone. Together, these results show that PFKL self-assembly is not a single pathway toward filaments, but a ligand-dependent supramolecular switching process in which distinct intermolecular interactions bias the architectural outcome.

## Results

### PFKL WT undergoes ligand-dependent switching between lattice-like and filamentous assemblies

PFKL WT forms micrometer-scale puncta in HepG2 cells, and its yeast homolog Pfk2p similarly assembles into punctate structures under hypoxic stress.^6,32^ However, conventional wide-field fluorescence microscopy is diffraction-limited and cannot determine whether these assemblies correspond to filaments, lattice-like structures, disordered clusters, or mixtures of distinct supramolecular states. To directly define how ligand state affects WT PFKL architecture, we used HS-AFM to monitor purified PFKL WT assembly in vitro under controlled ligand conditions. Because PFKL WT lattice-like assemblies were fragile under repeated HS-AFM scanning, we monitored lattice formation by imaging selected areas at defined time points for only a few frames, rather than continuously scanning the same lattice region.

Upon addition of 225 nM PFKL WT into 300 mM KCl buffer, HS-AFM imaging revealed spontaneous formation of two-dimensional lattice-like structures on mica under APO conditionss (**Fig. 2A** and **Fig. S2A**). These assemblies exhibited an average thickness of ∼10.2 nm, reduced long-range order, and heterogeneous surface topography with visible grain boundaries. Individual tetramers appeared densely packed but did not display filamentous morphologies, indicating that PFKL WT can form surface-associated lattice-like assemblies in the absence of filament-promoting ligand conditions.

**Figure 2.**
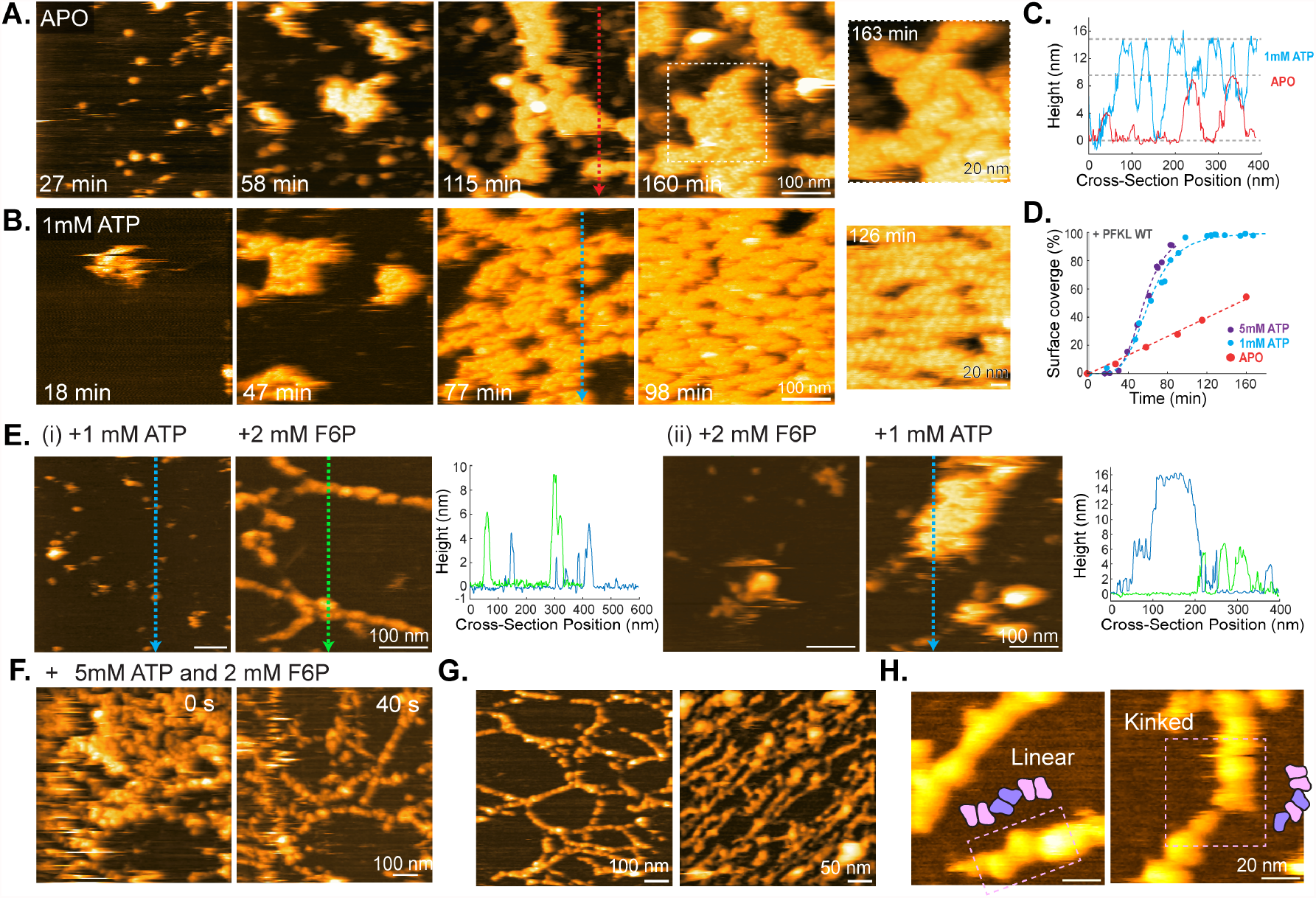
PFKL WT undergoes ligand-dependent switching between lattice-like and filamentous assemblies. **(A-B)** PFKL WT (225 nM) in buffer containing 300 mM KCl self-assembled into lattice-like structures on mica under **(A)** APO and **(B)** ATP conditions. **(C-D)** Cross-sectional height profiles and time-dependent surface coverage analysis of PFKL WT lattice growth under APO and ATP conditions. ATP promoted thicker, more ordered lattice-like islands and shifted lattice growth from approximately linear kinetics under APO conditionss to sigmoidal growth kinetics. Fitting parameters are summarized in **Table S3**; the 5 mM ATP dataset is shown in **Fig. S2C. (E)** Sequential HS-AFM measurements of PFKL WT self-assembly on mica under two treatment orders: **(i)** pre-incubation with 1 mM ATP followed by addition of 2 mM F6P, and **(ii)** pre-incubation with 2 mM F6P followed by addition of 1 mM ATP. ATP pre-binding followed by F6P addition redirected PFKL WT from lattice-like assembly toward filament formation (∼6.0 nm in thickness), whereas F6P-first treatment retained lattice-like organization. **(F)** Real-time HS-AFM imaging of PFKL WT filament formation in the presence of 5 mM ATP and 2 mM F6P. Filaments emerged beneath a partially dissociating lattice-like layer, supporting a ligand-dependent switch between structurally distinct assembly states. **(G-H)** Two distinct higher-order filament architectures were observed upon F6P addition: network-like and linear geometries. Zoomed-in views of linear and kinked filaments suggest that multiple tetramer-tetramer interfaces contribute to filament architecture.

We next tested whether ATP modulates WT assembly. In the presence of 1 mM or 5 mM ATP, PFKL WT formed more ordered lattice islands (**Fig. 2B** and **Fig. S2B-C**), accompanied by increased lattice thickness (**Fig. 2C**; ∼14.3 nm with ATP versus ∼9.6 nm in APO) and cooperative growth kinetics, as evidenced by sigmoidal growth curves for the analyzed lattice surface coverage (**Fig. 2D)**. Under APO conditionss, lattice expansion followed an approximately linear growth rate of 0.33% per min. In contrast, ATP promoted sigmoidal lattice growth curves with half-times of 60.9 min at 1 mM ATP and 55.9 min at 5 mM ATP, with Hill coefficients of 4.4 and 4.9, respectively (**Fig. 2D** and **Table S3**). The increased apparent height of ATP-associated assemblies suggests that ATP alters not only lattice-growth kinetics but also the structural organization of WT lattice-like assemblies. Thus, ATP does not induce filament formation on its own, but instead biases PFKL WT toward a cooperative lattice-prone assembly pathway. At 5 mM ATP, rapid lattice dissociation was observed during HS-AFM scanning (**Fig. S3**), suggesting that ATP enhances nucleation and cooperativity while forming a mature lattice that remains fragile under the applied HS-AFM scanning conditions.

We then asked whether substrate engagement redirects this ATP-supported lattice-like assembly pathway. Previous light scattering and electron microscopy (EM) studies reported dynamic filament formation of PFKL WT upon addition of F6P.^5,6^ To resolve the structural dynamics of this transition, we monitored ligand-dependent assembly using controlled, stepwise substrate addition during HS-AFM imaging.

Under APO or ATP conditions alone, PFKL WT assembled into 2D lattice-like structures (**Fig. 2A-D**). However, addition of F6P redirected the assembly outcome. When PFKL WT was pre-incubated with ATP and subsequently exposed to 2 mM F6P, filamentous structures ∼4-6 nm in thickness formed rapidly (**Fig. 2E-i** and **Fig. S4**). Time-lapse imaging revealed directional elongation, with individual PFKL units adding to filament ends, generating stable linear assemblies that persisted during continuous scanning (**Fig. S4B**). Thus, F6P does not simply enhance assembly; instead, in the context of ATP-prebound PFKL WT, it switches the supramolecular outcome from lattice-like assembly to filamentous structures.

The sequence of ligand engagement further influenced the structural trajectory. When F6P was added first and ATP was introduced afterward, PFKL WT retained the ∼15 nm lattice organization rather than converting into filaments (**Fig. 2E-ii**). Short incubations (∼20 min) with either ATP or F6P alone did not produce filamentous assemblies. These results indicate that filament formation requires coordinated ATP and F6P engagement and that ATP pre-binding prepares the tetramer for F6P-induced architectural switching.

At elevated ATP concentration (5 mM), F6P addition induced filament formation through local molecular rearrangement beneath a partially dissociating lattice layer (**Fig. 2F** and **Fig. S4C**), revealing that lattice and filament states are structurally distinct higher-order organizations rather than different stages of a single continuous assembly pathway. The transient coexistence of a dissociating lattice and emerging filaments further supports a ligand-driven switch in assembly state.

In addition to filament growth, two filament geometries were observed upon ATP and F6P treatment: linear and network-like assemblies (**Fig. 2G-H, Fig. S5**, and **Supplementary Movie 1**). Zoomed-in HS-AFM images revealed straight and kinked segments, suggesting that multiple tetramer-tetramer interfaces contribute to the filament architecture. These features are consistent with previous EM observations.^5^ Together, these observations establish that PFKL WT accesses distinct supramolecular states depending on ligand condition. APO and ATP-loaded states support lattice-like assemblies, whereas coordinated ATP and F6P engagement redirects PFKL WT toward filament formation. This ligand-order dependence suggests that substrate engagement does not simply increase or decrease self-assembly, but instead shifts the functional/conformational state of PFKL in a way that changes the intermolecular interactions available for higher-order organization. Thus, PFKL WT undergoes ligand-dependent supramolecular switching between lattice-like and filamentous assembly states.

### APO-PFKL WT forms lattice-like assemblies on continuous SLBs and membrane-patch substrates

The plasma membrane is a central hub of metabolic activity, supplying energy and biomass for cellular functions, such as actin rearrangement, in response to environmental cues.^22,33^ PFKL puncta have been observed localized near the plasma membrane in vivo.^6^ To test whether membrane association influences APO-PFKL WT higher-order organization, we reconstituted a supported lipid bilayer (SLB; DOPC/DOPS, 60:40) on mica and monitored APO-PFKL WT assembly by HS-AFM (**Fig. 3** and **Fig. S6**). Here, we used two membrane-supported substrates: continuous SLBs, which cover the mica surface and test whether APO-PFKL WT can assemble directly on a membrane, and membrane patches, which leave exposed mica gaps and membrane boundaries to compare where assembly initiates across different interfacial environments.

**Figure 3.**
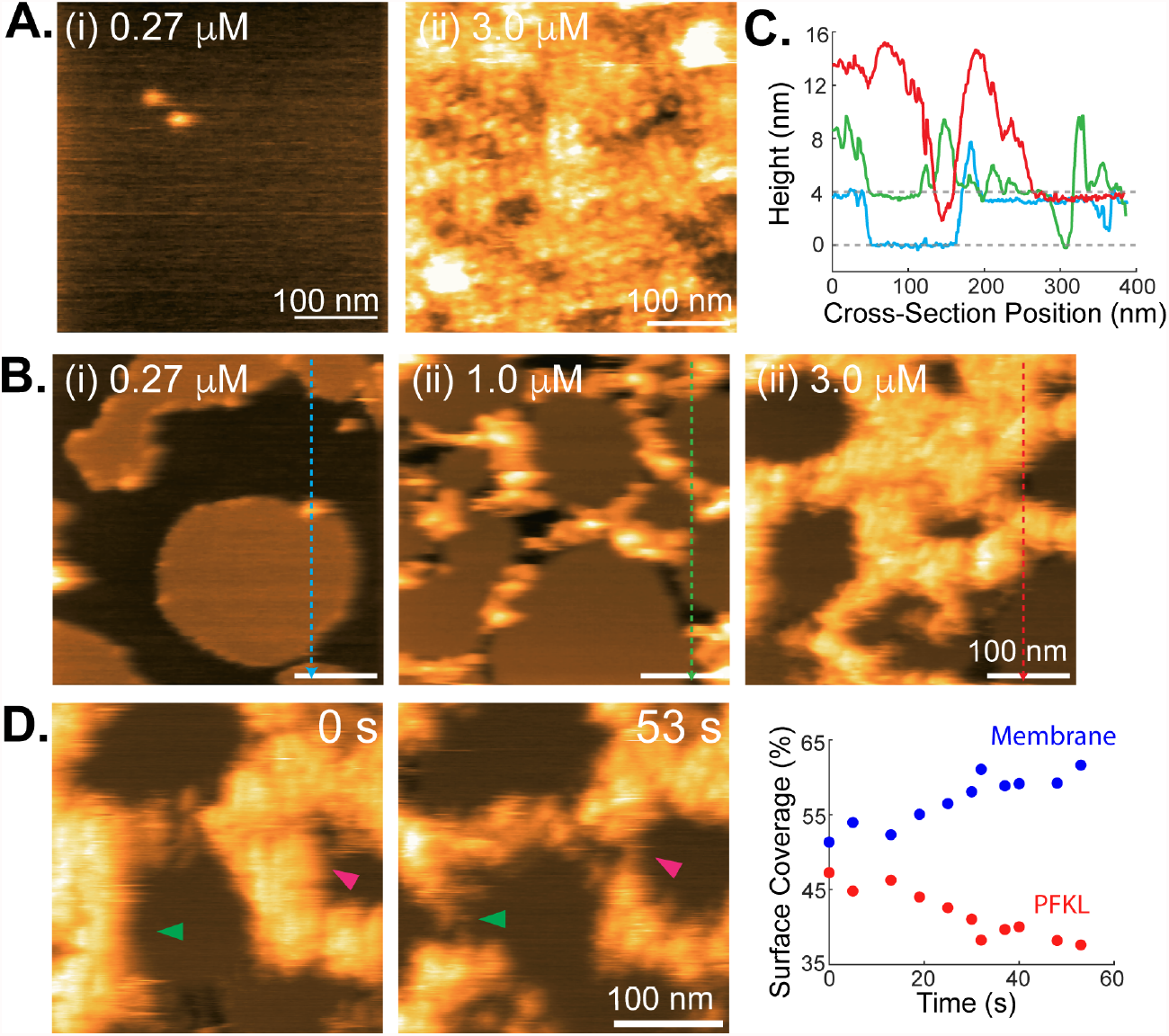
APO-PFKL WT self-assembles on continuous supported lipid bilayers and near membrane patches. **(A)** Representative HS-AFM images of PFKL WT bound to and assembled into lattice-like structures on a continuous, supported lipid bilayer composed of DOPC/DOPS (60:40). Bulk protein concentration modulates molecular binding density and lattice formation. **(B)** Representative HS-AFM images of PFKL WT self-assembled into lattice-like structures on mica containing pre-deposited membrane patches. **(C)** Cross-sectional height profiles measured across mica (∼0 nm), supported lipid bilayer (∼3.7 nm), and PFKL WT assemblies (∼9.5 nm and ∼13.2 nm). **(D)** Sequential HS-AFM images showing PFKL WT lattice-like assembly at exposed mica gaps between membrane patches and across membrane-supported regions. During scanning, the lattice gradually dissociated from the membrane (arrows).

On continuous SLBs, APO-PFKL WT formed lattice-like assemblies directly on the membrane surface under ligand-free conditions (**Fig. 3A** and **Fig. S6A-B**). To measure assembly height, we locally removed the protein layer by scratching a small region of the assembly after imaging and analyzed cross-sectional profiles relative to the exposed membrane surface. The membrane-supported assemblies showed an apparent height of ∼14 nm, similar to the ATP-associated lattice-like assemblies observed on mica (**Fig. 2B-C**). This indicates that APO-PFKL WT can adopt a comparable overall thickness on SLBs.

We next examined membrane patches pre-deposited on mica, which created exposed mica gaps between membrane patches. Under these heterogeneous surface conditions, lattice formation initiated predominantly on mica regions (**Fig. 3B, Fig. S6C-E**, and **Supplementary Movie 2**). As protein concentration increased, the lattices expanded laterally and extended across adjacent membrane patches. Height profiles again distinguished mica, membrane, and PFKL assemblies (**Fig. 3C**), confirming that the same lattice height ranges are preserved across both substrates.

Time-lapse HS-AFM imaging further revealed differential stability depending on the underlying support. Lattices formed at exposed mica gaps between membrane patches dissociated more rapidly during scanning than those associated with membrane patches (**Fig. 3D** and **Fig. S6F**). Together, these observations show that APO-PFKL WT can form lattice-like assemblies on or near membrane-supported surfaces. Continuous SLBs support lattice-like assembly, whereas membrane-patch substrates reveal that assembly initiation and scanning stability depend on the local interfacial environment. These results provide a membrane-relevant context for the WT lattice-like state before testing whether disruption of the previously described filament-forming interface alters access to lattice assembly.

### The filament-defective N702T mutant forms a cooperative double-layer lattice

Having established that PFKL WT forms lattice-like assemblies under APO and ATP conditions but switches to filaments upon coordinated ATP and F6P loading, we next asked whether disruption of the previously described N702-containing filament-forming interface abolishes higher-order assembly or instead reveals an alternative ordered assembly state. Because PFKL N702T is defective in filament formation^6^, this mutant provided a perturbation to separate filament formation from other potential modes of PFKL self-assembly.

To explore whether this mutant can self-assemble into higher-order structures, we first pre-incubated APO-PFKL N702T (100 nM in 100 mM KCl buffer) on mica for 10 min, and then imaged the assembled structures by HS-AFM (**Fig. 4A**). Notably, PFKL N702T formed highly ordered 2D lattices with clear periodicity, rather than filament-like structures. During HS-AFM imaging, the lattice gradually dissociated from its edge, and after ∼38 s, dissociation near the center exposed a second lattice layer (**Fig. 4A, Supplementary Movie 3**). Quantitative height analysis using cross-sectional profiles and a height histogram showed a total lattice thickness of ∼15.0 nm, consistent with a double-layer architecture of two ∼7.5 nm sheets (**Fig. 4B**, *top and middle panels*). Lattice surface coverage decayed exponentially during dissociation, with a halftime of 46.7 s (**Fig. 4B**, *bottom panel*; **Table S4**).

**Figure 4.**
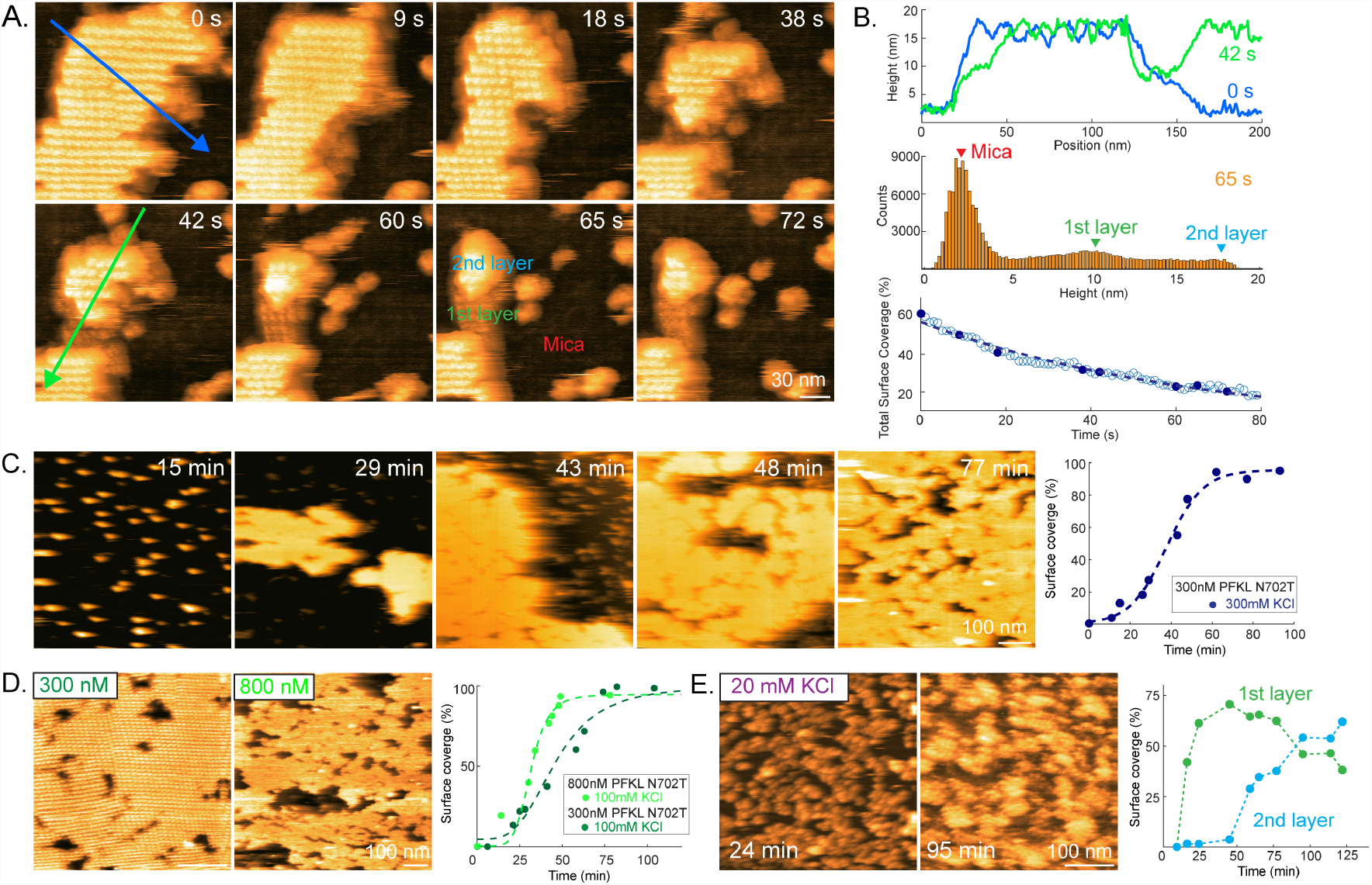
APO-PFKL N702T self-assembles into highly ordered double-layer 2D lattice. **(A)** HS-AFM images of the PFKL N702T 2D lattice formed by pre-incubating 100 nM protein on mica for 10 minutes in buffer containing 100 mM KCl. During imaging, the lattice dissociates from the edge, revealing a two-layer architecture. **(B)** Cross-section height profiles, height histogram, and surface-coverage analysis of lattice dissociation. **(C)** Time-lapse HS-AFM showing lattice assembly at 300 nM PFKL N702T in 300 mM KCl across different scan areas. **(D)** Lattice formation at 300 nM and 800 nM PFKL N702T in 100 mM KCl. Representative end-point images are shown on the left, and time-dependent surface coverage fitted with sigmoidal curves is shown on the right. Fitting parameters, including *t*_1/2_ and *n*_*H*_, are summarized in **Table S3. (E)** At low ionic strength, 300 nM PFKL N702T forms island-like assemblies with locally ordered lattice regions. Time-dependent lattice surface coverage is shown on the right.

To identify the biochemical conditions that mediate lattice formation, we varied protein concentration (100-500 nM), incubation time (3-15 min), and KCl concentration (20-300 mM). Ordered double-layer lattices were consistently observed at 100-300 mM KCl, whereas lowering KCl to 20 mM disrupted long-range lattice order (**Fig. S7**). At 20 mM KCl, PFKL N702T assemblies appeared disorganized but retained local periodicity, indicating that reduced ionic strength permits local contacts but is insufficient to support long-range lattice organization. Across all HS-AFM observations, larger continuous lattices were stable during HS-AFM scanning, whereas small lattice patches were fragile, indicating that lattice continuity affects imaging stability rather than lattice formation itself.

Next, to visualize the self-assembly dynamics of fragile lattices, APO-PFKL N702T (300 nM) was directly supplemented into the liquid cell, and newly scanned regions containing lattice assemblies formed at different time points were analyzed prior to dissociation. This experimental strategy enabled kinetic analysis of lattice formation before scanning-induced disruption occurred. PFKL N702T molecules first deposited as small aggregates, followed by the appearance of ∼15.0 nm-thick lattice islands after ∼29 min (**Fig. 4C**). These lattice islands expanded laterally and fused into continuous lattices with visible grain boundaries. The characteristic surface coverage followed sigmoidal growth kinetics with a growth half-time of 36.3 min and a Hill coefficient of 5.0 (**Fig. 4C** and **Table S3**), indicating cooperative assembly of the lattice state^34,35^. We noted that no additional layers formed after the double-layer lattice appeared, consistent with surface-associated nucleation followed by lateral growth at the solid-liquid interface^27,29^.

We then examined ionic strength dependence of PFKL N702T lattice formation at 20 mM and 100 mM KCl (**Fig. 4D-E, Fig. S8**, and **Table S3**). Under the same protein concentration (300 nM), increasing KCl from 100 to 300 mM accelerated lattice assembly (**Fig. 4C-D, Supplementary Movie 4 and 5, and Table S3;** *t*_1/2_ decreased from 46.1 to 36.3 min; *n*_*H*_ increased from 3.5 to 5.0). The distinct *n*_*H*_ value reflects that PFKL N702T self-assembly at 300 mM KCl has a steeper rise during the rapid growth phase of lattice formation. At constant ionic strength (100 mM KCl), increasing protein concentration from 300 to 800 nM further accelerated lattice growth (**Fig. 4D and Table S3;** *t*_1/2_ decreased from 46.1 to 31.6 min; *n*_*H*_ increased from 3.5 to 6.1). At low KCl concentration, assembly followed a different pathway: a single layer formed first, and a second layer subsequently stacked on top (**Fig. 4E, Supplementary Movie 6**). The resulting structures lacked long-range periodic order despite local lattice-like domains (**Fig. S8D-i**).

These results support cooperative nucleation followed by lateral expansion rather than sequential tetramer addition as the primary lattice-growth model. The persistence of ordered lattice assembly in the filament-defective mutant PFKL N702T further indicates that the lattice does not arise from disrupted filament polymerization but instead represents a distinct assembly pathway. In addition, the sensitivity of lattice organization to ionic conditions suggests that the biochemical environment biases the structural route of PFKL self-organization rather than merely modulating assembly stability. These observations establish the double-layer 2D lattice as an alternative higher-order assembly state accessible to PFKL.

### PFKL N702T tetramers adopt a defined orientation within the double-layer lattice

To determine how PFKL N702T tetramers are organized within the double-layer lattice, we quantitatively defined the geometric parameters of individual lattice units under distinct biochemical conditions. Because PFKL WT undergoes ligand-dependent switching from lattice-like assemblies under APO/ATP conditions to filaments upon ATP/F6P loading, we next asked whether N702T preserves lattice organization under the same ligand conditions. PFKL N702T (100 nM) was incubated on freshly cleaved mica in 100 mM KCl buffer to allow rapid lattice formation and then imaged under APO, ATP, and ATP+F6P conditions (**Fig. 5A-B, Supplementary Movie 7**). Across all three states, PFKL N702T formed highly ordered lattices with indistinguishable packing symmetry, adopting a hexagonal arrangement with a centered motif. Thus, N702T preserves the same lattice geometry across ligand conditions, uncoupling lattice formation from the ATP/F6P-induced filament transition observed for PFKL WT.

**Figure 5.**
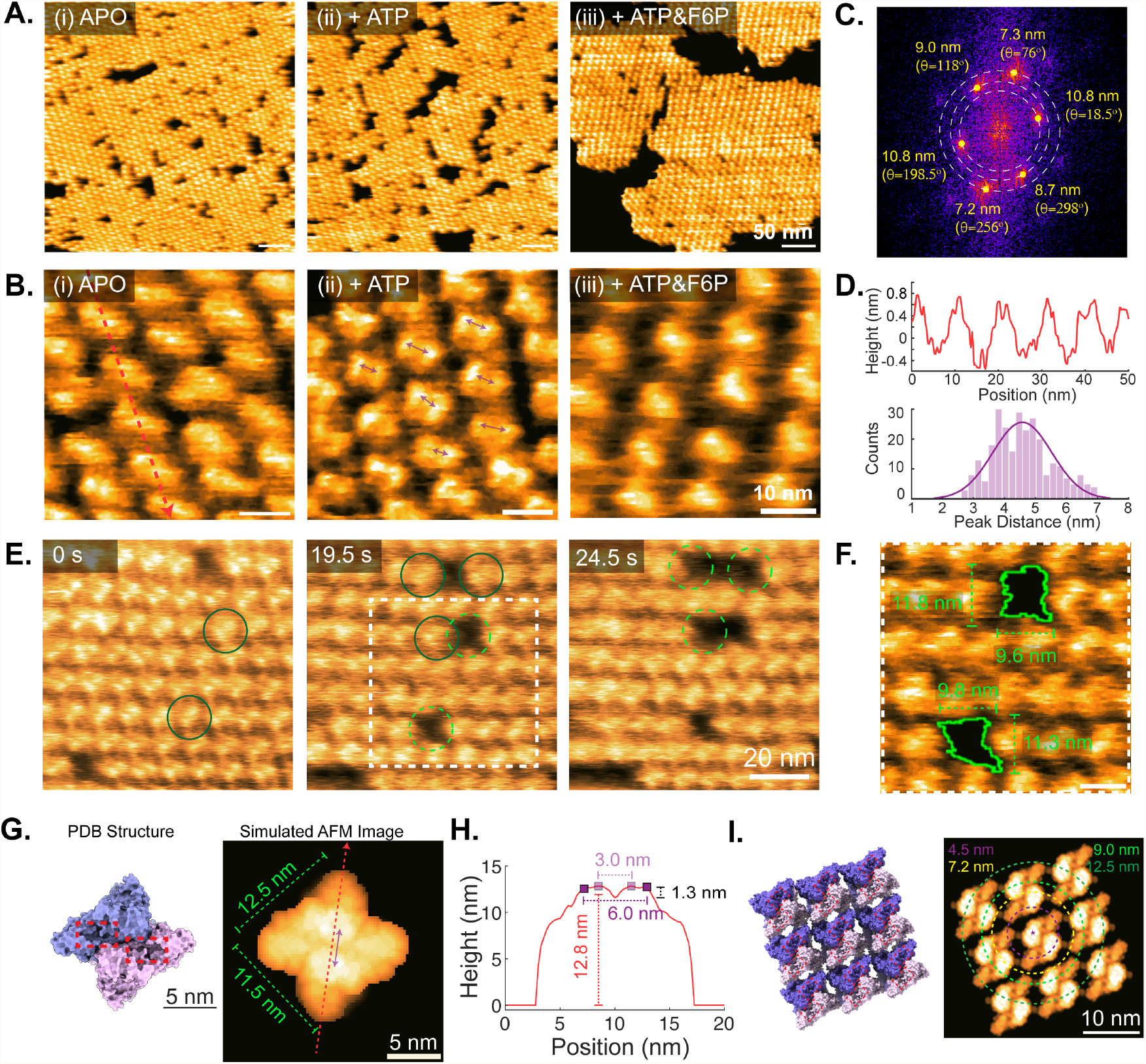
Orientation of PFKL N702T tetramers within the 2D lattice. **(A-B)** Representative HS-AFM images of PFKL N702T lattices formed under APO, ATP, and ATP+F6P conditions at different length scales. **(C)** Masked FFT image of the PFKL N702T lattice shown in **(A-i)** under APO conditions, revealing the hexagonal diffraction peaks. **(D)** *Top:* Cross-sectional height profile of the lattice shown in **(B-i**) under APO conditions. *Bottom:* Histogram of measured distances between the two protruding height maxima within a lattice unit (purple arrows in **B-ii**, ATP condition), fitted with a Gaussian distribution yielding 4.5 ± 1.0 nm. **(E-F)** Sequential dissociation of PFKL N702T molecules from the lattice. **(G-H)** Simulated AFM image of PFKL tetramer (PDB: 8W2G; tip radius: ∼1 nm) viewed from the dimeric cap orientation, showing two active sites within the catalytic domains (red rectangles). Two protruding height maxima are separated by 3-6 nm with an inter-peak depth of ∼1.3 nm. **(I)** Simulated AFM image of a PFKL lattice composed of nine tetramers, with circular radii overlaid to match experimental spacing.

To define lattice symmetry, we performed spatial FFT analysis on the APO lattice acquired over a large scan area (**Fig. 5A-(i)**). The FFT map displayed distinct hexagonal peaks corresponding to intermolecular spacings of ∼7.2 nm, ∼8.8 nm, and ∼10.8 nm at defined angular orientations (**Fig. 5C**). This spatial periodicity confirms a long-range hexagonally ordered 2D arrangement as shown in high-resolution HS-AFM images (**Fig. 5B, Supplementary Movie 8**). Cross-sectional height profiles of the lattice (**Fig. 5D**, *top*) revealed periodic protrusions with an average amplitude of ∼0.8 nm and lateral spacing of ∼11.0 nm. The histogram of distances between the two protruding height maxima within each lattice unit (**Fig. 5D**, *bottom*) followed a Gaussian distribution centered at 4.5 ± 1.0 nm. These measurements provide a reproducible topographic fingerprint of the repeating lattice unit.

To directly relate lattice geometry to tetramer architecture, we analyzed sequential dissociation events (**Fig. 5E-F, Supplementary Movie 9**). Vacant lattice sites generated during molecular dissociation exhibited projected dimensions of ∼9.7 × 11.5 nm. These dimensions define the in-lattice footprint of individual tetramers and indicate a constrained orientation distinct from that expected for freely adsorbed tetramers. We next simulated AFM images derived from the cryo-EM structure of the PFKL tetramer using a 1 nm probe model, and then compared the resulting spatial features with these experimental observations (**Fig. 5G-H** and **Fig. S9**). By systematically comparing rotational orientations, we identified a configuration in which the dimeric cap containing the catalytic domains faces the cantilever with a defined tilt generated by a 90-degree rotation about the short two-fold axis. This orientation quantitatively reproduces the measured lateral footprint (**Fig. 5G**; ∼12.5 × 11.5 nm), the inter-peak spacing of 3-6 nm, and the inter-peak depth (**Fig. 5H**; ∼1.3 nm) observed experimentally. Simulated multi-tetramer lattices constructed using this orientation reproduced the periodic features seen in HS-AFM images (**Fig. 5I**), further supporting the assignment.

These experimental observations and structural analyses establish that PFKL N702T tetramers assemble into a hexagonally packed double-layer lattice. The catalytic dimeric cap is consistently exposed to solution in a defined orientation, fixing the spatial presentation of active-site-containing domains while constraining intermolecular interfaces. Thus, the lattice represents a structurally organized state with defined architectural rules rather than a nonspecific aggregate^26,27^. Importantly, the preservation of this orientation across ligand conditions demonstrates that lattice geometry is structurally stable and decoupled from catalytic-state switching. These results establish a direct structural link between tetramer architecture and higher-order organization, providing a structural basis for treating the lattice as a genuine supramolecular state rather than a nonspecific or collapsed assembly.

### MD simulations identify distributed inter-tetramer contacts stabilizing the N702T lattice

To determine how PFKL N702T forms and stabilizes lattices in the absence of ATP and F6P, we constructed a seven-tetramer lattice model based on the APO geometry observed by HS-AFM and performed all-atom MD simulations (**Fig. 6A**). The lattice remained structurally intact throughout the trajectory, despite anisotropic motion of the peripheral tetramers relative to the central tetramer, indicating that the APO lattice is flexible but stable as a higher-order assembly (**Fig. 6B**).

**Figure 6.**
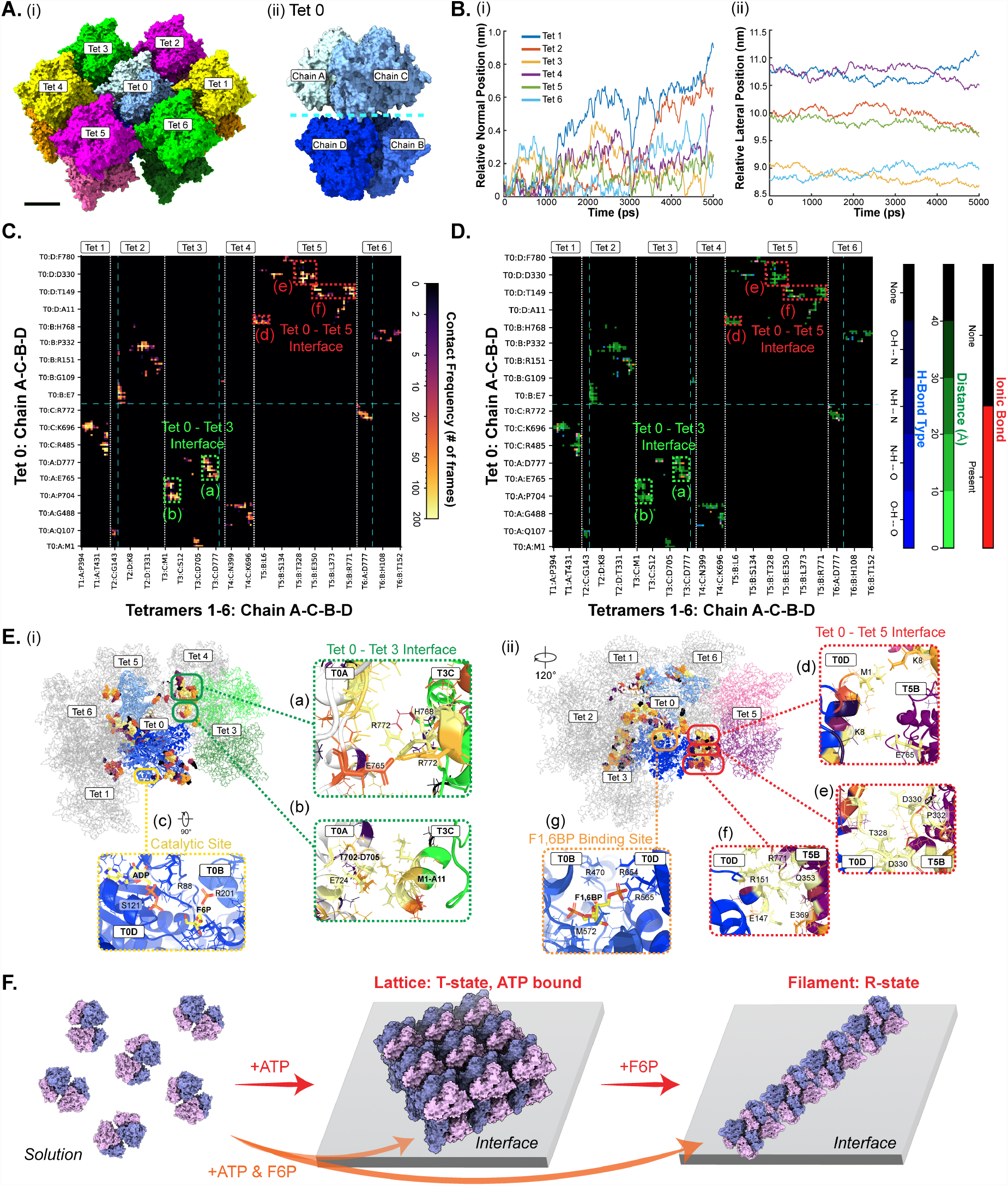
MD simulations define inter-tetramer contacts in the PFKL N702T lattice. **(A)** Surface representation of the initial PFKL N702T lattice assembly composed of seven tetramers, constructed based on the lattice organization inferred from HS-AFM data (**Fig. 5B-i and Fig. 5I**). Equivalent tetramer-tetramer interfaces are indicated by matching colors. **(B)** Time-dependent positions of each exterior tetramer relative to the central tetramer in a representative all-atom MD simulation trajectory. **(C-D)** Residue-residue contact maps showing **(C)** the frequency of inter-tetramer contacts, and **(D)** a multichannel contact map of the inter-tetramer interaction network in the PFKL N702T lattice structure. Only residues participating in persistent inter-tetramer contacts are shown; additional details are provided in **Table S7** and **Fig. S10**. The white dotted lines and cyan dashed lines define the boundaries between neighboring tetramers and between chains AC and BD, respectively. **(E)** Representative interfacial contacts in the PFKL N702T lattice structure, with residues making contacts colored according to contact frequency and shown as spheres. Tet 2 in (i) and Tet 4 in (ii) are omitted for clarity. Zoomed-in views highlight two representative inter-tetramer interfaces, together with the catalytic site and ligand-binding site in the central tetramer. F6P, F1,6BP, and ADP were superimposed onto the simulated MD lattice structure using PDB structure 8W2G. **(F)** Proposed working model for ligand-dependent supramolecular switching of PFKL WT. APO-PFKL WT forms a thinner lattice-like assembly, whereas ATP-bound PFKL WT forms a thicker 2D double-layer lattice-like assembly. Upon coordinated ATP and F6P loading, lattice-like and filamentous assemblies coexist, consistent with F6P-driven redirection toward filament formation. Interface regions indicate sites where PFKL tetramers engage neighboring tetramers to initiate, stabilize, or remodel supramolecular assemblies.

Interfacial residue contact-map analysis over the 2-4 ns interval of the MD trajectory showed that the lattice is supported by a distributed inter-tetramer interaction network comprising potential electrostatic interactions, hydrogen-bonding interactions and loop-mediated contacts across multiple neighboring interfaces, rather than by a single dominant residue pair (**Fig. 6C-D** and **Table S7**). The most persistent contacts were observed at representative interfaces between T0A-T3C and T0D-T5B (**Fig. 6E**). Here, the first index denotes the tetramer and the second denotes the chain.

At representative interfaces, the inter-tetramer contact network involved multiple structural regions rather than a single localized binding surface. For example, the T0A-T3C interface included contacts between C-terminal segments of adjacent chains, as well as contacts between the Thr702-Asp705 loop on T0A and the N-terminal helix of T3C (**Fig. 6E-i-a,b**). A second representative interface, T0D-T5B, involved contacts between the N-terminal region of one chain and the C-terminal region of the neighboring tetramer, together with loop-mediated interactions between internal and C-terminal segments (**Fig. 6E-i-d,e**). These examples illustrate that lattice stabilization arises from a distributed set of contacts across several regions of the tetramer, including loop-loop, hydrophobic, electrostatic, and hydrogen-bonding interactions. Additional representative interfacial contacts are shown in **Table S7** and **Fig. S10**.

The T0A-T3C interface notably includes the N702T mutation site, which occupies a peripheral position, contacts Val4 and Glu7 on adjacent subunits with only moderate occupancy (**Fig. 6E-i-b**), and did not form a Thr702-Thr702 interaction via hydrogen bonding. The other three T702 residues in the central tetramer also do not participate in close inter-tetramer contacts. Thus, residue 702 is not the principal determinant of lattice formation, even though it may contribute to local stabilization. Together with the HS-AFM observation that N702T forms ordered lattices rather than filaments, this result supports the conclusion that lattice assembly uses intermolecular contacts distinct from the previously described N702-dependent filament interface.

This structural organization also suggests how ligand binding may influence assembly selection. The mapped F6P- and ADP-binding regions remain exposed to solution (**Fig. 6E-i-c**), suggesting that the lattice architecture is compatible with substrate access. By contrast, the mapped F1,6BP-binding site lies near an inter-tetramer interface (**Fig. 6E-i-g**). This arrangement suggests that ligand binding could differentially affect higher-order assembly: whereas access to the catalytic site is preserved in the lattice, occupancy of the F1,6BP-binding region may influence the geometry or stability of specific inter-tetramer contacts. In this framework, the APO lattice remains compatible with substrate access, while substrate-dependent shifts in interface selection may favor the alternative filament-promoting assembly mode in PFKL WT.

Because lattice stabilization is mediated by distributed contacts across multiple interfaces rather than by a single dominant interaction, local dissociation does not require failure of the entire assembly. In this view, the HS-AFM observation of dissociation from lattice edges is consistent with disruption of a subset of the inter-tetramer contacts that maintain the local lattice environment, while preserving the overall lattice architecture. These simulations support a model in which the APO PFKL N702T lattice is stabilized by a distributed lateral interaction network that is structurally distinct from the interface associated with PFKL filament formation. This model provides a framework for understanding why N702T preserves lattice assembly while PFKL WT can be redirected toward filament formation by coordinated ATP and F6P loading.

## Discussion

PFK-1 is a highly regulated, rate-controlling enzyme in central carbon metabolism that modulates glycolytic flux through ligand-dependent stabilization of inactive and active conformational states. Our findings show that, for the liver isoform PFKL, this regulation extends beyond intratetramer allostery to ligand-dependent switching between structurally distinct higher-order assembly states. Rather than representing nonspecific clustering, the assemblies observed here reflect supramolecular states whose formation depends on ligand conditions and intermolecular interactions between tetramers.

The central finding of this study is that PFKL WT does not follow a single assembly pathway toward filaments. Instead, PFKL WT forms lattice-like assemblies under APO and ATP conditions, whereas coordinated ATP and F6P loading redirects the enzyme toward filament formation (**Fig. 2**). This ligand-order dependence suggests that substrate engagement shifts the functional or conformational state of PFKL in a way that changes the intermolecular interactions available for higher-order organization. Thus, ligand binding does not merely tune the extent of self-assembly; it changes the architectural outcome of PFKL self-organization. The height difference between APO and ATP-associated PFKL WT lattice-like assemblies further suggests that ligand binding may alter not only assembly kinetics but also the preferred packing geometry of WT tetramers within lattice-like states (**Fig. 2A-C**). By analogy to the structurally defined N702T lattice, the thinner APO assemblies may reflect a more tilted or laterally adsorbed arrangement, whereas the thicker ATP-associated assemblies may approach a more upright or layered packing geometry. Because WT lattice-like assemblies are less ordered and less stable during HS-AFM imaging than the N702T lattice, these assignments should be considered putative packing models rather than definitive structural orientations.

The N702T mutant provides a mechanistic perturbation that separates lattice formation from filament formation. Although N702T disrupts the previously described filament-forming interface, it does not abolish higher-order assembly. Instead, N702T forms a highly ordered double-layer lattice under APO conditionss and preserves this lattice geometry under ATP and ATP+F6P conditions (**Figs. 4-5**). Together with the WT data, these results indicate that lattice formation is not unique to the mutant, but that N702T stabilizes a more ordered and reproducible version of a lattice-like state accessible to PFKL WT. Importantly, unlike PFKL WT, N702T fails to undergo the ATP/F6P-induced transition to filaments. This contrast suggests that residue 702 is not required for lattice formation itself, but contributes to the ligand-dependent transition from lattice-like assembly to filament formation.

The combined HS-AFM, AFM image simulation, and MD analyses support a model in which the N702T lattice is a defined supramolecular architecture rather than a defective filament or nonspecific aggregate. HS-AFM resolves the surface topography of the lattice, and comparison with topology-based AFM image simulations defines the molecular packing geometry of PFKL tetramers within the lattice (**Fig. 5**). MD simulations further suggest that the lattice is stabilized by distributed contacts across multiple interfaces, including representative contacts at the T0A-T3C and T0D-T5B interfaces (**Fig. 6**). This distributed-interface model also helps explain the ionic-strength dependence of lattice formation: ordered lattices formed at 100-300 mM KCl, whereas 20 mM KCl supported only local periodicity without long-range order (**Fig. 4**). Because the MD model identifies electrostatic, hydrogen-bonding, hydrophobic, and loop-mediated contacts across neighboring tetramers, changes in ionic strength may alter the balance of interactions required for local contacts to mature into extended lattice order.

The MD model further suggests how ligand binding could influence assembly selection. Specifically, the mapped F6P- and ADP-binding regions remain exposed to solution, suggesting that the lattice architecture is compatible with substrate access, whereas the mapped F1,6BP-binding site lies near an inter-tetramer interface (**Fig. 6E**). This arrangement indicates that ligand binding may differentially affect lattice and filament assembly by altering the geometry or stability of specific inter-tetramer contacts. In this framework, the APO lattice remains compatible with substrate access, while substrate-dependent shifts in interface selection may favor the filament-promoting assembly mode in PFKL WT.

Together, these observations support a working model in which PFKL WT lattice-like assemblies are dynamic, ligand-responsive states rather than static scaffolds (**Fig. 6F**). The coexistence of ordered lattice structure with local dissociation suggests that PFKL assemblies are dynamically maintained rather than statically locked (**Fig. 2** and **Fig. 4**). Under APO or ATP-loaded conditions, PFKL WT can form lattice-like assemblies, and elevated ATP promotes cooperative lattice growth. Because ATP functions both as a substrate and as a concentration-dependent allosteric regulator of PFKL^6^, the thicker ATP-associated lattice-like state may represent a ligand-stabilized organization in which PFKL is spatially organized but not yet committed to filament formation. Sequential addition of F6P to ATP-bound PFKL WT induces filament formation, whereas simultaneous addition of ATP and F6P leads to coexistence of lattice-like and filamentous assemblies. In this working model, both lattice-like and filamentous assemblies may represent functional supramolecular states, but they differ in stability and ligand dependence. Lattice-like assemblies provide one ligand-stabilized organizational mode, whereas filaments represent a substrate-loaded assembly mode that appears more stable during ATP+F6P-induced coexistence and may support localized glycolytic organization or recruitment to specific cellular regions. Direct catalytic consequences of lattice and filament states remain to be tested, but the present data link ligand sensing, conformational state, and supramolecular organization.

Our membrane-reconstitution experiments place the WT lattice-like state in a membrane-relevant context. APO-PFKL WT forms lattice-like assemblies on or near membrane-supported surfaces, and membrane-patch substrates reveal that assembly initiation and scanning stability depend on the local interfacial environment (**Fig. 3** and **Fig. S6**). This membrane-relevant context is important because glycolytic enzymes often localize near the plasma membrane, where glucose uptake occurs through GLUT transporters and where localized ATP production can support cytoskeletal remodeling, cell migration, and other processes with high local energy demand.^21,36,37^ In this context, PFKL supramolecular switching may provide a structural mechanism for organizing glycolytic activity in spatially defined cellular regions, including glycolysis-dependent states such as cancer. ^21,37^

More broadly, our findings support a model in which PFKL self-assembly functions as a tunable mesoscale layer of glycolytic regulation. By shifting between lattice-like and filamentous architectures, PFKL may alter its structural context and ligand responsiveness on second-to-minute timescales, depending on protein concentration and ligand condition. This work supports the view that metabolic control can emerge not only from active-site chemistry and classical allostery, but also from reversible changes in supramolecular organization.

## ASSOCIATED CONTENT

### Supporting Information

Materials and methods, supporting figures and tables, and supporting movie captions.

## AUTHOR INFORMATION

### Author contributions

S.L. and Y.-C.L. designed research. S.L. performed HS-AFM experiments. A.P. performed MD simulations under the supervision of Y.-C.L. N.P.-E. developed an AFM simulator based on protein PDB structure. S.L., A.P. and Y.-C.L. analyzed the data and wrote the manuscript. B.A.W. prepared the protein samples. All authors reviewed and edited the manuscript.

## Acknowledgements

This work was supported by start-up funds from the University of Texas at Austin and the U.S. National Institutes of Health grant R35 GM150528 to S.L. and Y.-C.L., U.S. National Institutes of Health grant R35 GM158392 to B.A.W., U.S. National Institutes of Health grant R01 AI169412, R01 GM144472, R01 DA043571, and the Welch Foundation F-2143 to K.-L.H. Heather Hansen (WVU) is thanked for her technical and intellectual contributions. The authors acknowledge the Texas Advanced Computing Center (TACC) at The University of Texas at Austin for providing computational resources that have contributed to the research results reported within this paper. URL: http://www.tacc.utexas.edu

## Competing interests

The authors declare no competing interests.

### Data availability

The manuscript figures, supplementary figures, and supplementary movies contain all data necessary to interpret, verify and extend the presented work. The raw data files can be obtained from the authors upon reasonable request.

