## Supplemental Information for "High-Speed AFM Reveals Ligand-Dependent Supramolecular Switching of Human Phosphofructokinase-1"

### Experimental Methods:

#### Human PFKL protein and cryo-EM structures

Recombinant N-terminal His-tagged human PFKL WT (NP_002617) and PFKL N702T, were purified from Sf9 cells as reported previously.[1] The cryo-EM structure of R-state PFKL tetramer (PDB: 8W2G) and T-state PFKL tetramer (PDB: 8W2H) were collected from Protein Data Bank and illustrated by UCSF ChimeraX.[2-4]

#### HS-AFM Experiments

The basic imaging buffer consisted of 50 mM HEPES (pH 7.4), 2 mM $CaCl_2$, and 10 mM $MgCl_2$, supplemented with the indicated KCl concentration (100 mM or 300 mM). PFKL protein samples were either deposited onto freshly cleaved mica or introduced directly into the fluid cell of the HS-AFM for in situ assembly experiments.

All movies were acquired in amplitude modulation mode using a high-speed AFM (MS-NEX, RIBM Co., Ltd., Japan). Ultrashort cantilevers (NanoWorld, Switzerland) with spring constants of 0.15 or 0.6 $N \cdot m^{-1}$, resonance frequency of 0.6 MHz, and a quality factor of ~1.5 in buffer were used. Cantilevers were sharpened by oxygen plasma etching if needed. Tip–sample interactions were minimized by maintaining the set-point/free-amplitude ratio at ~0.85, with a free amplitude of ~1.0 nm. HS-AFM images were recorded at 1-5 frames/s.

#### HS-AFM Image Processing and Data Analysis

All HS-AFM images were processed in ImageJ[5]. Surface coverage and height histograms from HS-AFM frames were analyzed using ImageJ. Further quantitative analyses, including sigmoidal growth fitting and dissociation kinetics, were analyzed by custom routines in MATLAB R2024a. Simulated AFM images (4 pixels/nm resolution) were generated using a custom Python code by convolving the protein PDB structure with a cone-shape tip (tip radius 1 nm, cone angle 15°).

#### MD Simulations

The starting coordinates for all-atom molecular dynamics (MD) simulations were derived from the cryo-EM structure of the wild-type PFKL tetramer in the T-state (PDB: 8W2H)[1], with missing N-terminal residues 1-12 in each chain modeled using AlphaFold[6] and the N702T mutation performed with the PyMOL mutagenesis tool[7]. In this study, only APO structure was simulated, without ligands. To create the lattice model, six surrounding tetramers were positioned relative to the central tetramer to reproduce the observed lattice symmetry in HS-AFM (**Fig. 5C**). The initial edge-to-edge distance between adjacent tetramers was set to a minimum of 0.3 nm to avoid trapping the simulation in a local energy minimum caused by strong hydrogen bond formation. Throughout the analysis, tetramer nomenclature is as follows: T0 (or Tet 0) is the central tetramer, surrounded by 6 surrounding tetramers (Tet 1-Tet 6 or T1-T6). Tetramers 1 and 4, 2 and 5, and 3 and 6 were initially symmetry-related. Chains were named alphabetically within each tetramer, such that the first residue in the sequence would be Tetramer 0 Chain A Met 1 (T0A:M1).

Simulations were performed using the GROMACS software package (version 2024.2)[8]. The CHARMM36[9] force field was used to describe the protein and ions, and water molecules were modeled using the TIP3P model, which is the water model for which the CHARMM36 force field is parameterized[10]. The system was neutralized and solvated in a 150 mM KCl aqueous solution to mimic the imaging conditions and placed in a triclinic simulation box expanded to ensure a minimum distance of 2 nm between periodic images of protein molecules. The resulting box was composed of 3,307,506 atoms and enclosed a volume of $3.3 \times 10^4$ $nm^3$ (**Table S5**)

All simulations were performed using the leap-frog integrator[11] with a time-step of 2 fs. Bonds involving hydrogen atoms were constrained using the LINCS algorithm[12]. Short-range electrostatic and van der Waals interactions were truncated at 1.0 nm using a Verlet cutoff scheme. Long-range electrostatic interactions were computed using the particle mesh Ewald

(PME) method[13]. Long-range dispersion corrections were applied to both energy and pressure. Periodic boundary conditions were applied in all three dimensions.

Energy minimization was performed using the steepest descent algorithm in two stages. In the first stage, water molecules were allowed to relax while harmonic positional restraints were applied to protein heavy atoms. Following this, protein atoms were allowed to relax in the second stage to decrease the maximum force below 1000 kJ mol$^{-1}$ nm$^{-1}$. Following minimization, the system was equilibrated using a multi-step process in which the force constants of harmonic position constraints were gradually decreased from 1000 kJ mol$^{-1}$ nm$^{-2}$ to 0 kJ mol$^{-1}$ nm$^{-2}$ to allow relaxation of the protein lattice within the solvent environment. The first two steps were performed in the NVT ensemble, while the latter four steps were performed in the NPT ensemble. More details regarding the system setup and equilibration process can be found in **Table S6**.

During production simulations, the temperature of the system was maintained at 310 K using the velocity-rescaling thermostat[14] with a time constant of 0.1 ps. Temperature coupling was applied separately to the protein and solvent components of the system. Pressure was maintained at 1.0 bar using the stochastic cell-rescaling barostat[15] with isotropic coupling and a time constant of 5 ps. A single production trajectory of 10 ns was generated. Atom positions and energies were saved every 10 ps. Prior to analysis, trajectories were processed in GROMACS to remove periodic boundary condition artifacts and then aligned translationally and rotationally to the initial structure. Structural analyses were performed using the MDAnalysis python package (version 2.10)[16,17]. The equilibrium portion of the trajectory, determined from stabilization of center-of-mass distances between neighboring tetramers and the central tetramer, as well as the RMSD from the initial post-equilibrium structure, corresponding to t = 2-4 ns of simulation time, was isolated for analysis. The lattice plane was defined based on the centers-of-mass of neighboring tetramers and recomputed on a per-frame basis to measure displacement in the lateral and normal directions relative to the plane. The plane was fit to the array of points with singular-value decomposition (SVD).

The potential for inter-protein contacts was predicted based on a threshold criterion of Cβ distances within 1.0 nm of each other[18]. The capability of electrostatic interactions was predicted on a per-residue basis by determining whether a residue pair had opposite charges. The potential for a hydrogen bond was predicted solely based on hydrogen bond donor-acceptor identities present in side chains. For further analysis, residues forming a persistent contact in each simulation were defined as being within the threshold distance for at least 25% of the surveyed frames.

All production simulations were performed on the Texas Advanced Computing Center (TACC) Lonestar6 high-performance computing cluster.

### Supplementary Tables:

#### Table S1. HS-AFM Experimental Conditions for PFKL WT

| Figures | Dynamics | Molecular players | | | Imaging buffer | | | Protein Deposition | |
|---|---|---|---|---|---|---|---|---|---|
| | | WT (nM) | F6P (mM) | ATP (mM) | KCl (mM) | $Mg^{2+}$ (mM) | $Ca^{2+}$ (mM) | Mica incubation (min) | Live addition |
| 2A | Lattice-like Assembly | 225 | | | 300 | 10 | 2 | | V |
| 2B | Lattice-like Assembly | 225 | | 1 | 300 | 10 | 2 | | V |
| 2E | (i) Filament Assembly | 255 | 2 (2nd) | 1(1st) | 300 | 10 | 2 | | V |
| | (ii) Lattice-like Assembly | 170 | 2 (1st) | 1 (2nd) | 300 | 10 | 2 | | V |
| 2F | Filament Assembly | 225 | 2 | 5 | 300 | 10 | 2 | | V |
| 2G | (left) Network Geometry | 255 | 2 | 1 | 300 | 10 | 2 | | V |
| | (right) Linear Geometry | 425 | 2 | 1 | 300 | 10 | 2 | | V |
| 2H | Linearized Filament | 425 | 2 | 1 | 300 | 10 | 2 | | V |
| | Kinked Filament | 425 | 2 | 1 | 300 | 10 | 2 | | V |
| S2A | Self-Assembly | 225 | | | 300 | 10 | 2 | | V |
| S2B | Self-Assembly | 225 | | 1 | 300 | 10 | 2 | | V |
| S2C | Self-Assembly | 225 | | 5 | 300 | 10 | 2 | | V |
| S3A | Disassociation | 225 | | 5 | 300 | 10 | 2 | | V |
| S4A | (i) Filament Assembly | 170 | 2 (2nd) | 1 (1st) | 300 | 10 | 2 | | V |
| | (ii) Filament Assembly | 425 | 2 (2nd) | 1 (1st) | 300 | 10 | 2 | | V |
| S4B | Filament Assembly | 170 | 2 (2nd) | 1 (1st) | 300 | 10 | 2 | | V |
| S4C | Filament Assembly | 225 | 2 | 5 | 300 | 10 | 2 | | V |
| S5A | Filament Assembly (Network) | 255 | 2 | 1 | 300 | 10 | 2 | | V |
| S5B | Filament Assembly (Linear) | 425 | 2 | 1 | 300 | 10 | 2 | | V |
| 3A | Single Molecular Characterization | 225 | | | 100 | 10 | 2 | | V |
| 3B | Membrane Interaction | 270 | | | 100 | 10 | 2 | 5 | |
| | | 1000 | | | 100 | 10 | 2 | 3 | |
| | | 3000 | | | 100 | 10 | 2 | 5 | |
| 3D | Membrane Interaction | 3000 | | | 100 | 10 | 2 | 5 | |

**Table S1. HS-AFM Experimental Conditions for PFKL WT (continued)**

| Figures | Dynamics | Molecular players | | | Imaging buffer | | | Protein Deposition | |
|---|---|---|---|---|---|---|---|---|---|
| | | WT (nM) | F6P (mM) | ATP (mM) | KCl (mM) | $Mg^{2+}$ (mM) | $Ca^{2+}$ (mM) | Mica incubation (min) | Live addition |
| S6A | Single Molecular Characterization | 225→270 | | | 100 | 10 | 2 | | V |
| S6B | Membrane Interaction | 2700 | | | 100 | 10 | 2 | 1 | |
| S6C-D | Membrane Interaction | 1000 | | | 100 | 10 | 2 | 3 | |
| S6E-F | Membrane Interaction | 3000 | | | 100 | 10 | 2 | 5 | |

**Table S2. HS-AFM Experimental Conditions for PFKL N702T**

| Figures | Dynamics | Molecular players | | | Imaging buffer | | | Protein Deposition | |
|---|---|---|---|---|---|---|---|---|---|
| | | N702T (nM) | F6P (mM) | ATP (mM) | KCl (mM) | $Mg^{2+}$ (mM) | $Ca^{2+}$ (mM) | Mica incubation (min) | Live addition |
| 4A | Lattice Dissociation | 100 | | | 100 | | 2 | 10 | |
| 4C* | Self-Assembly | 300 | | | 300 | | 2 | | V |
| 4D | Self-Assembly | 300 | | | 100 | | 2 | | V |
| | | 800 | | | 100 | | 2 | | V |
| | | 300 | | | 300 | | 2 | | V |
| 4E | Self-Assembly | 300 | | | 20 | | 2 | | V |
| 5A-B | (i) 2D Lattice Formation | 90 | | | 300 | 10 | 2 | | V |
| 5A-B | (ii) 2D Lattice Formation | 90 | | 1 | 300 | 10 | 2 | | V |
| 5A-B | (iii) 2D Lattice Formation | 170 | 2 | 1 | 300 | 10 | 2 | | V |
| 5E | Lattice Dissociation | 100 | | | 100 | | 2 | 10 | |
| S7A | Self-Assembly | 100 | | | 100 | | 2 | 3 | |
| | | 100 | | | 100 | | 2 | 10 | |
| | | 500 | | | 100 | | 2 | 5 | |
| | | 500 | | | 100 | | 2 | 10 | |
| | | 500 | | | 100 | | 2 | 15 | |
| | | 300 | | | 20 | | 2 | 10 | |
| | | 300 | | | 300 | | 2 | 10 | |
| S8A | Self-Assembly | 300 | | | 20 | | 2 | | V |
| S8B | (i) Self-Assembly | 800 | | | 100 | | 2 | | V |
| | (ii) Self-Assembly | 300 | | | 100 | | 2 | | V |
| S8C | Self-Assembly | 300 | | | 300 | | 2 | | V |
| S8D | (i) Self-Assembly | 300 | | | 20 | | 2 | | V |
| | (ii) Self-Assembly | 300 | | | 100 | | 2 | | V |

*: After full lattice coverage had formed on mica, 2 mM F6P was added at 77 min, followed by 1 mM ATP at 93 min.

**Table S3. Summary of Sigmoidal Fitting Parameters for Lattice Growth**

| Figures | Dynamics | PFKL | Functional state | Experiment condition | | Baseline (%) | Hill Coefficient | Half Time ($t_{1/2}$, min) | Top (%) |
|---|---|---|---|---|---|---|---|---|---|
| | | | | Protein (nM) | KCl (mM) | | | | |
| 2D | Self-Assembly | WT | APO | 225 | 300 | N.A. (linear fit) | | | |
| | | WT | +1 mM ATP | 225 | 300 | 0 | 4.4 | 60.9 | 100 |
| | | WT | +5 mM ATP | 225 | 300 | 0 | 4.9 | 55.9 | 100 |
| 4C* | Self-Assembly | N702T | APO | 300 | 300 | 3.8 | 5.0 | 36.3 | 95.8 |
| 4D | Self-Assembly | N702T | APO | 300 | 100 | 4.7 | 3.5 | 46.1 | 100 |
| | | N702T | APO | 800 | 100 | 0.4 | 6.1 | 31.6 | 94.5 |

*: During the final stage of imaging, 2 mM F6P (77 min) and 1 mM ATP (93 min) were sequentially added.

**Table S4. PFKL Lattice Dissociation Kinetics**

| Figures | Dynamics | PFKL | Experiment conditions: Protein (nM) | Experiment conditions: KCl (mM) | Amplitude | k | Half Time |
|---|---|---|---|---|---|---|---|
| S3B | Dissociation (5 mM ATP) | WT | 225 | 300 | 77.00% | 0.1957 $s^{-1}$ | 3.5 s |
| 4B (bottom) | Dissociation | N702T | 100 | 100 | 57.13% | 0.0148 $s^{-1}$ | 46.7 s |

**Table S5. Experimental Setups for Molecular Dynamics (MD) simulations**

| System (PFKL N702T) | Box Volume ($nm^3$) | Total Atoms | Protein Atoms | Water Molecules | KCl concentration (mM) |
|---|---|---|---|---|---|
| PFKL N702T Lattice (Seven Tetramers) | 33415.2 | 3307506 | 334180 | 989758 | 150 |

**Table S6. Equilibration Protocol for MD Simulations**

| Step | Ensemble | Length (ps) | Time Step (fs) | Harmonic Restraint Force Constant (kJ $mol^{-1}$ $nm^{-2}$) |
|---|---|---|---|---|
| **1** | NVT | 100 | 2 | 1000 |
| **2** | NVT | 100 | 2 | 250 |
| **3** | NPT | 100 | 1 | 250 |
| **4** | NPT | 200 | 1 | 50 |
| **5** | NPT | 400 | 2 | 10 |
| **6** | NPT | 100 | 2 | 0 |

### Table S7. Inter-tetramer Interactions with Central Dimer in MD Simulation (from 2 ns to 4 ns)

| Edge Tetramer (Tetramer-Chain) | | Central Tetramer (Tetramer-Chain) | | Contact Type | | Contact Frequency | Distance (Å) | |
|---|---|---|---|---|---|---|---|---|
| Chain ID | Residue | Chain ID | Residue | Electrostatic | Hydrogen Bond | | Mean | SD |
| T1A | LYS395 | T0C | GLY697 | FALSE | FALSE | 59 | 8.94 | 0.63 |
| T1A | GLU396 | T0C | LYS696 | TRUE | N-H..O | 150 | 7.38 | 1.38 |
| T1A | GLU396 | T0C | GLY697 | FALSE | FALSE | 184 | 7.39 | 1.03 |
| T1A | LYS397 | T0C | LYS696 | FALSE | FALSE | 142 | 7.33 | 1.61 |
| T1A | LYS397 | T0C | GLY697 | FALSE | FALSE | 125 | 7.65 | 2 |
| T1A | LYS397 | T0C | ARG698 | FALSE | FALSE | 59 | 8.01 | 0.94 |
| T1A | SER398 | T0C | LYS696 | FALSE | N-H..O | 93 | 8.56 | 1.23 |
| T1A | SER398 | T0C | GLY697 | FALSE | FALSE | 154 | 8.23 | 1.08 |
| T1A | SER398 | T0C | ARG698 | FALSE | FALSE | 77 | 9.21 | 0.55 |
| T1A | ASN399 | T0C | GLU515 | FALSE | N-H..O | 58 | 9.44 | 0.41 |
| T1A | ASN399 | T0C | LYS696 | FALSE | N-H..O | 145 | 7.6 | 1.3 |
| T1A | ASN399 | T0C | GLY697 | FALSE | FALSE | 127 | 7.52 | 1.27 |
| T1A | ASN399 | T0C | ARG698 | FALSE | FALSE | 169 | 8.24 | 0.98 |
| T1A | TYR694 | T0C | ARG485 | FALSE | FALSE | 76 | 9.34 | 0.49 |
| T1A | ARG695 | T0C | GLU482 | TRUE | FALSE | 89 | 9.2 | 0.59 |
| T1A | ARG695 | T0C | ARG485 | FALSE | FALSE | 116 | 8.89 | 0.64 |
| T1A | LYS696 | T0C | GLU478 | TRUE | N-H..O | 167 | 8.97 | 0.72 |
| T1A | LYS696 | T0C | VAL481 | FALSE | FALSE | 127 | 9.04 | 0.48 |
| T1A | LYS696 | T0C | GLU482 | TRUE | N-H..O | 198 | 7.62 | 0.72 |
| T1A | LYS696 | T0C | ARG485 | FALSE | FALSE | 167 | 8.43 | 1.01 |
| T1A | GLY697 | T0C | VAL481 | FALSE | FALSE | 150 | 9.19 | 0.46 |
| T1A | GLY697 | T0C | GLU482 | FALSE | FALSE | 146 | 8.89 | 0.66 |
| T1A | GLY697 | T0C | ARG485 | FALSE | FALSE | 196 | 7.41 | 1.05 |
| T1A | GLY697 | T0C | TYR514 | FALSE | FALSE | 50 | 9.48 | 0.41 |
| T1A | GLY697 | T0C | GLU515 | FALSE | FALSE | 136 | 8.61 | 0.86 |
| T1A | GLY697 | T0C | GLU516 | FALSE | FALSE | 154 | 8.78 | 0.58 |
| T2C | GLY143 | T0A | GLY62 | FALSE | FALSE | 52 | 8.76 | 0.95 |
| T2D | MET1 | T0B | ASP5 | FALSE | FALSE | 106 | 8.33 | 1.08 |
| T2D | MET1 | T0B | LYS8 | FALSE | FALSE | 179 | 7.57 | 1.44 |
| T2D | MET1 | T0B | LEU9 | FALSE | FALSE | 91 | 8.46 | 0.91 |
| T2D | MET1 | T0B | SER12 | FALSE | FALSE | 90 | 8.05 | 1.25 |
| T2D | ALA2 | T0B | ASP5 | FALSE | FALSE | 71 | 8.18 | 1.06 |
| T2D | ALA2 | T0B | LYS8 | FALSE | FALSE | 80 | 7.73 | 1.55 |
| T2D | GLU326 | T0B | ASP330 | FALSE | FALSE | 108 | 8.42 | 0.94 |
| T2D | ALA327 | T0B | ASP330 | FALSE | FALSE | 81 | 9.22 | 0.55 |
| T2D | THR328 | T0B | THR328 | FALSE | FALSE | 69 | 9.06 | 0.63 |
| T2D | THR328 | T0B | PRO329 | FALSE | FALSE | 115 | 9.13 | 0.66 |
| T2D | THR328 | T0B | ASP330 | FALSE | FALSE | 196 | 6.48 | 1.56 |
| T2D | THR328 | T0B | THR331 | FALSE | FALSE | 114 | 8.56 | 1.08 |
| T2D | THR328 | T0B | PRO332 | FALSE | FALSE | 50 | 9.31 | 0.52 |
| T2D | PRO329 | T0B | ASP330 | FALSE | FALSE | 115 | 9.06 | 0.66 |
| T2D | ASP330 | T0B | ASP330 | FALSE | FALSE | 116 | 7.59 | 1.26 |
| T2D | ASP330 | T0B | THR331 | FALSE | FALSE | 50 | 9.19 | 0.59 |
| T2D | ASP330 | T0B | PRO332 | FALSE | FALSE | 61 | 8.93 | 0.63 |
| T2D | THR331 | T0B | ASP330 | FALSE | FALSE | 118 | 7.86 | 1.12 |
| T2D | GLU767 | T0B | ARG151 | TRUE | FALSE | 65 | 9.42 | 0.48 |
| T2D | HIS768 | T0B | GLU137 | TRUE | N-H..O | 65 | 9.27 | 0.52 |
| T2D | THR770 | T0B | GLU147 | FALSE | FALSE | 53 | 9.33 | 0.5 |
| T2D | ARG772 | T0B | GLU147 | TRUE | FALSE | 69 | 8.76 | 0.93 |
| T3C | MET1 | T0A | GLU724 | FALSE | FALSE | 148 | 8.45 | 0.9 |
| T3C | MET1 | T0A | LYS727 | FALSE | FALSE | 95 | 9.15 | 0.59 |
| T3C | VAL4 | T0A | THR702 | FALSE | FALSE | 67 | 9.16 | 0.61 |
| T3C | VAL4 | T0A | ALA703 | FALSE | FALSE | 186 | 8.21 | 1.05 |
| T3C | VAL4 | T0A | PRO704 | FALSE | FALSE | 200 | 6.26 | 0.49 |
| T3C | VAL4 | T0A | ASP705 | FALSE | FALSE | 91 | 9.06 | 0.61 |
| T3C | VAL4 | T0A | PRO721 | FALSE | FALSE | 160 | 9.32 | 0.43 |
| T3C | VAL4 | T0A | THR723 | FALSE | FALSE | 194 | 8.35 | 0.7 |
| T3C | VAL4 | T0A | GLU724 | FALSE | FALSE | 170 | 8.9 | 0.63 |
| T3C | ASP5 | T0A | PRO704 | FALSE | FALSE | 110 | 8.88 | 0.63 |
| T3C | ASP5 | T0A | GLU724 | FALSE | FALSE | 80 | 9.29 | 0.34 |

| | | | | | | | | |
|---|---|---|---|---|---|---|---|---|
| T3C | GLU7 | T0A | THR702 | FALSE | FALSE | 54 | 9.34 | 0.5 |
| T3C | GLU7 | T0A | ALA703 | FALSE | FALSE | 129 | 8.97 | 0.63 |
| T3C | GLU7 | T0A | PRO704 | FALSE | FALSE | 85 | 9.44 | 0.43 |
| T3C | LYS8 | T0A | ALA703 | FALSE | FALSE | 163 | 8.49 | 0.98 |
| T3C | LYS8 | T0A | PRO704 | FALSE | FALSE | 114 | 8.76 | 0.74 |
| T3C | LYS8 | T0A | ASP705 | TRUE | N-H..O | 85 | 9.48 | 0.36 |
| T3C | ALA11 | T0A | ALA703 | FALSE | FALSE | 89 | 9.13 | 0.57 |
| T3C | GLN372 | T0A | HIS768 | FALSE | N-H..N | 71 | 9.03 | 0.6 |
| T3C | ALA703 | T0A | VAL4 | FALSE | FALSE | 56 | 9.13 | 0.72 |
| T3C | PRO704 | T0A | MET1 | FALSE | FALSE | 80 | 8.62 | 0.96 |
| T3C | PRO704 | T0A | ALA3 | FALSE | FALSE | 77 | 8.87 | 0.69 |
| T3C | PRO704 | T0A | VAL4 | FALSE | FALSE | 57 | 9.14 | 0.64 |
| T3C | ASP705 | T0A | MET1 | FALSE | FALSE | 96 | 8.7 | 0.84 |
| T3C | ASP705 | T0A | VAL4 | FALSE | FALSE | 84 | 8.6 | 0.58 |
| T3C | PRO721 | T0A | MET1 | FALSE | FALSE | 61 | 9.24 | 0.5 |
| T3C | GLU724 | T0A | MET1 | FALSE | FALSE | 52 | 9.2 | 0.53 |
| T3C | HIS768 | T0A | ARG772 | FALSE | FALSE | 138 | 9.38 | 0.41 |
| T3C | HIS768 | T0A | ASP777 | TRUE | N-H..O | 161 | 8.42 | 0.68 |
| T3C | HIS768 | T0A | LYS778 | FALSE | N-H..N | 82 | 8.83 | 0.81 |
| T3C | HIS768 | T0A | GLY779 | FALSE | FALSE | 111 | 8.57 | 0.99 |
| T3C | THR770 | T0A | THR770 | FALSE | FALSE | 183 | 8.04 | 0.76 |
| T3C | THR770 | T0A | ARG771 | FALSE | FALSE | 130 | 8.41 | 1.01 |
| T3C | THR770 | T0A | ARG772 | FALSE | FALSE | 175 | 7.69 | 1.08 |
| T3C | THR770 | T0A | ASP777 | FALSE | FALSE | 134 | 7.88 | 1.11 |
| T3C | ARG771 | T0A | GLU767 | TRUE | FALSE | 113 | 9.38 | 0.43 |
| T3C | ARG771 | T0A | HIS768 | FALSE | FALSE | 145 | 9.11 | 0.47 |
| T3C | ARG771 | T0A | THR770 | FALSE | FALSE | 147 | 7.64 | 1.31 |
| T3C | ARG771 | T0A | ARG771 | FALSE | FALSE | 70 | 8.86 | 0.59 |
| T3C | ARG771 | T0A | ARG772 | FALSE | FALSE | 88 | 8.99 | 0.52 |
| T3C | ARG772 | T0A | THR770 | FALSE | FALSE | 87 | 9.26 | 0.58 |
| T3C | ARG772 | T0A | ARG772 | FALSE | FALSE | 85 | 9.07 | 0.6 |
| T3C | ASP777 | T0A | GLU767 | FALSE | FALSE | 67 | 8.37 | 0.5 |
| T3C | LYS778 | T0A | GLU767 | TRUE | N-H..O | 56 | 9.33 | 0.52 |
| T4C | ILE486 | T0A | LYS696 | FALSE | FALSE | 125 | 9.07 | 0.55 |
| T4C | GLY488 | T0A | LYS696 | FALSE | FALSE | 110 | 8.87 | 0.78 |
| T4C | GLY488 | T0A | GLY697 | FALSE | FALSE | 128 | 8.95 | 0.72 |
| T4C | LYS696 | T0A | ILE486 | FALSE | FALSE | 81 | 8.19 | 1.64 |
| T4C | LYS696 | T0A | GLY488 | FALSE | FALSE | 89 | 8.79 | 0.78 |
| T4C | GLY697 | T0A | ASN399 | FALSE | FALSE | 112 | 9.03 | 0.7 |
| T4C | GLY697 | T0A | GLY488 | FALSE | FALSE | 92 | 8.74 | 1.02 |
| T5B | MET1 | T0D | MET1 | FALSE | FALSE | 107 | 7.81 | 1.1 |
| T5B | MET1 | T0D | ALA2 | FALSE | FALSE | 57 | 7.13 | 2.02 |
| T5B | MET1 | T0D | VAL4 | FALSE | FALSE | 68 | 8.28 | 1.13 |
| T5B | ALA2 | T0D | MET1 | FALSE | FALSE | 124 | 8.47 | 0.95 |
| T5B | ALA2 | T0D | ALA2 | FALSE | FALSE | 55 | 8.78 | 1.18 |
| T5B | ASP5 | T0D | MET1 | FALSE | FALSE | 143 | 7.15 | 1.47 |
| T5B | ASP5 | T0D | ALA2 | FALSE | FALSE | 58 | 8.02 | 1.39 |
| T5B | LYS8 | T0D | MET1 | FALSE | FALSE | 72 | 8.54 | 0.9 |
| T5B | LEU9 | T0D | MET1 | FALSE | FALSE | 56 | 8.05 | 1.17 |
| T5B | ARG129 | T0D | ASP330 | TRUE | FALSE | 91 | 9.57 | 0.34 |
| T5B | SER130 | T0D | PRO329 | FALSE | FALSE | 76 | 9.41 | 0.43 |
| T5B | SER130 | T0D | ASP330 | FALSE | O-H..O | 200 | 7.22 | 0.73 |
| T5B | GLY133 | T0D | ASP330 | FALSE | FALSE | 85 | 9.2 | 0.53 |
| T5B | HIS155 | T0D | ASP330 | TRUE | N-H..O | 70 | 9.31 | 0.44 |
| T5B | PRO329 | T0D | THR328 | FALSE | FALSE | 142 | 9.18 | 0.51 |
| T5B | PRO329 | T0D | ASP330 | FALSE | FALSE | 87 | 9.29 | 0.5 |
| T5B | ASP330 | T0D | GLU326 | FALSE | FALSE | 200 | 6.82 | 0.73 |
| T5B | ASP330 | T0D | ALA327 | FALSE | FALSE | 193 | 8.85 | 0.49 |
| T5B | ASP330 | T0D | THR328 | FALSE | FALSE | 200 | 5.76 | 0.59 |
| T5B | ASP330 | T0D | ASP330 | FALSE | FALSE | 174 | 8.56 | 0.86 |
| T5B | ASP330 | T0D | THR331 | FALSE | FALSE | 198 | 7.84 | 0.65 |
| T5B | ASP330 | T0D | LEU766 | FALSE | FALSE | 128 | 9.29 | 0.51 |
| T5B | THR331 | T0D | THR328 | FALSE | FALSE | 187 | 8.85 | 0.4 |
| T5B | PRO332 | T0D | THR328 | FALSE | FALSE | 200 | 8.02 | 0.75 |
| T5B | PRO332 | T0D | PRO329 | FALSE | FALSE | 148 | 8.96 | 0.78 |
| T5B | PRO332 | T0D | ASP330 | FALSE | FALSE | 71 | 8.79 | 0.81 |

| | | | | | | | | |
|---|---|---|---|---|---|---|---|---|
| T5B | MET349 | T0D | PRO329 | FALSE | FALSE | 176 | 8.72 | 0.65 |
| T5B | MET349 | T0D | ASP330 | FALSE | FALSE | 63 | 9.55 | 0.36 |
| T5B | GLU350 | T0D | ARG151 | TRUE | FALSE | 200 | 8.41 | 0.57 |
| T5B | GLU350 | T0D | SER154 | FALSE | O-H..O | 98 | 9.32 | 0.43 |
| T5B | GLN353 | T0D | ARG151 | FALSE | FALSE | 200 | 7.62 | 0.46 |
| T5B | GLN353 | T0D | SER154 | FALSE | O-H..O | 142 | 9.39 | 0.36 |
| T5B | MET354 | T0D | THR148 | FALSE | FALSE | 67 | 9.73 | 0.22 |
| T5B | MET354 | T0D | ARG151 | FALSE | FALSE | 196 | 8.62 | 0.53 |
| T5B | GLU357 | T0D | GLU147 | FALSE | FALSE | 185 | 8.64 | 0.66 |
| T5B | GLU357 | T0D | THR148 | FALSE | FALSE | 66 | 9.64 | 0.27 |
| T5B | GLU357 | T0D | ARG151 | TRUE | FALSE | 165 | 9.15 | 0.46 |
| T5B | LYS360 | T0D | GLU147 | TRUE | N-H..O | 66 | 9.49 | 0.4 |
| T5B | GLU369 | T0D | GLU147 | FALSE | FALSE | 61 | 9.34 | 0.4 |
| T5B | GLU765 | T0D | LYS8 | TRUE | N-H..O | 158 | 9.43 | 0.41 |
| T5B | HIS768 | T0D | ALA11 | FALSE | FALSE | 125 | 8.73 | 0.91 |
| T5B | HIS768 | T0D | ALA14 | FALSE | FALSE | 112 | 8.89 | 0.85 |
| T5B | HIS768 | T0D | GLY15 | FALSE | FALSE | 108 | 9.21 | 0.57 |
| T5B | VAL769 | T0D | ARG151 | FALSE | FALSE | 101 | 9.4 | 0.51 |
| T5B | VAL769 | T0D | THR152 | FALSE | FALSE | 90 | 9.24 | 0.6 |
| T5B | ARG771 | T0D | THR148 | FALSE | FALSE | 200 | 7.45 | 0.63 |
| T5B | ARG771 | T0D | ARG151 | FALSE | FALSE | 51 | 9.62 | 0.29 |
| T5B | ARG771 | T0D | THR152 | FALSE | FALSE | 174 | 8.96 | 0.63 |
| T5B | LEU774 | T0D | THR148 | FALSE | FALSE | 166 | 9.01 | 0.64 |
| T5B | LEU774 | T0D | ARG151 | FALSE | FALSE | 136 | 9.13 | 0.54 |
| T5B | MET776 | T0D | SER146 | FALSE | FALSE | 53 | 9.53 | 0.41 |
| T5B | MET776 | T0D | GLU147 | FALSE | FALSE | 84 | 9.54 | 0.32 |
| T5B | MET776 | T0D | THR148 | FALSE | FALSE | 200 | 6.37 | 0.71 |
| T5B | MET776 | T0D | ARG151 | FALSE | FALSE | 124 | 9.18 | 0.53 |
| T5B | MET776 | T0D | THR152 | FALSE | FALSE | 63 | 9.56 | 0.31 |
| T5B | ASP777 | T0D | THR148 | FALSE | FALSE | 138 | 9.26 | 0.44 |
| T6A | THR770 | T0C | THR770 | FALSE | FALSE | 64 | 9.08 | 0.66 |
| T6A | THR770 | T0C | ARG771 | FALSE | FALSE | 94 | 8.95 | 0.7 |
| T6A | ARG771 | T0C | THR770 | FALSE | FALSE | 58 | 8.73 | 0.63 |
| T6A | ARG771 | T0C | ARG771 | FALSE | FALSE | 53 | 9.02 | 0.56 |
| T6A | ARG772 | T0C | THR770 | FALSE | FALSE | 69 | 8.62 | 0.86 |
| T6A | ASP777 | T0C | HIS768 | TRUE | N-H..O | 148 | 8.89 | 0.68 |
| T6A | ASP777 | T0C | THR770 | FALSE | FALSE | 62 | 7.98 | 0.86 |
| T6A | LYS778 | T0C | HIS768 | FALSE | N-H..N | 175 | 7.77 | 1.7 |
| T6A | LYS778 | T0C | THR770 | FALSE | FALSE | 51 | 8.6 | 0.4 |
| T6A | GLY779 | T0C | HIS768 | FALSE | FALSE | 50 | 8.77 | 0.64 |
| T6A | PHE780 | T0C | HIS768 | FALSE | FALSE | 90 | 8.28 | 1.27 |

Note: The colored zones highlight spatial contacts within local interfaces between adjacent tetramers in the MD simulations. Each color corresponds to a zoomed-in structural view shown in **Fig. 6** and **Fig. S10**.

### Supplementary Figures:

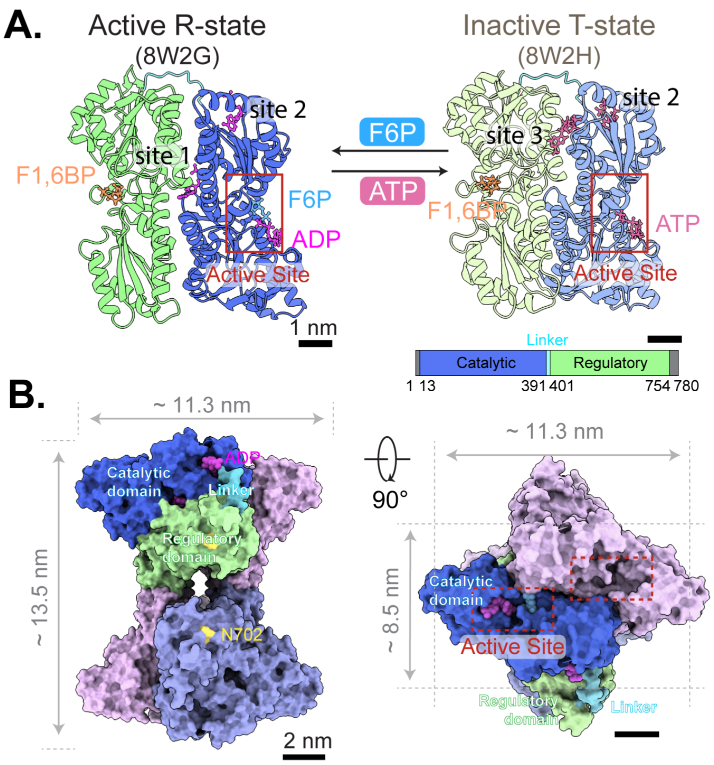

**Figure S1. Architectures of PFKL from monomeric form to tetramer. (A)** Monomeric structures of PFKL in its active R-state (PDB: 8W2G; with F1,6BP, F6P, and ADP) and inactive T-state (PDB: 8W2H; with F1,6BP and ATP). PFKL is involved in catalyzing the phosphorylation of F6P to F1,6BP with ATP hydrolysis at active site (red rectangle). **(B)** PFKL tetramer in the active R-state (PDB: 8W2G). With 90 degrees of rotation, the extrusions of PFKL tetramer show a dimension of ~11.3 nm and ~8.5 nm, with two active sites. Within one of the monomers, the catalytic domain (dark blue), interdomain linker (light blue), and regulatory domain (purple) are highlighted, respectively. The N702 residue (yellow) is crucial to form filamentous structures.

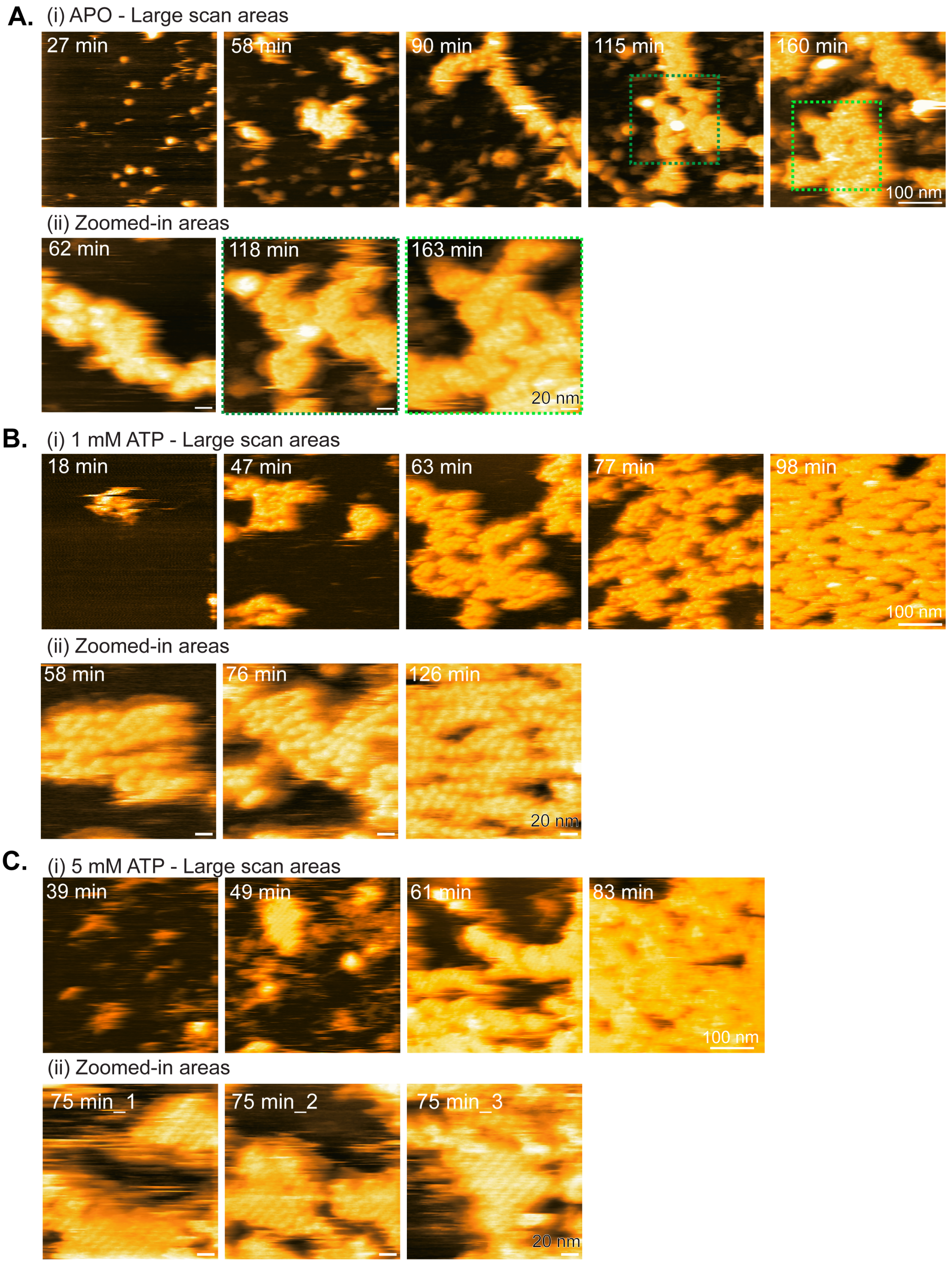

**Figure S2. Time-lapse HS-AFM imaging of PFKL WT lattice-like assembly under APO and ATP conditions. (A-C)** Representative HS-AFM images showing PFKL WT (225 nM) self-assembling into lattice-like assemblies in buffer containing 300 mM KCl under APO, 1 mM ATP, or 5 mM ATP conditions. Lattice-like domains progressively expand over time, with ATP-treated conditions showing more cooperative growth of ordered two-dimensional assemblies.

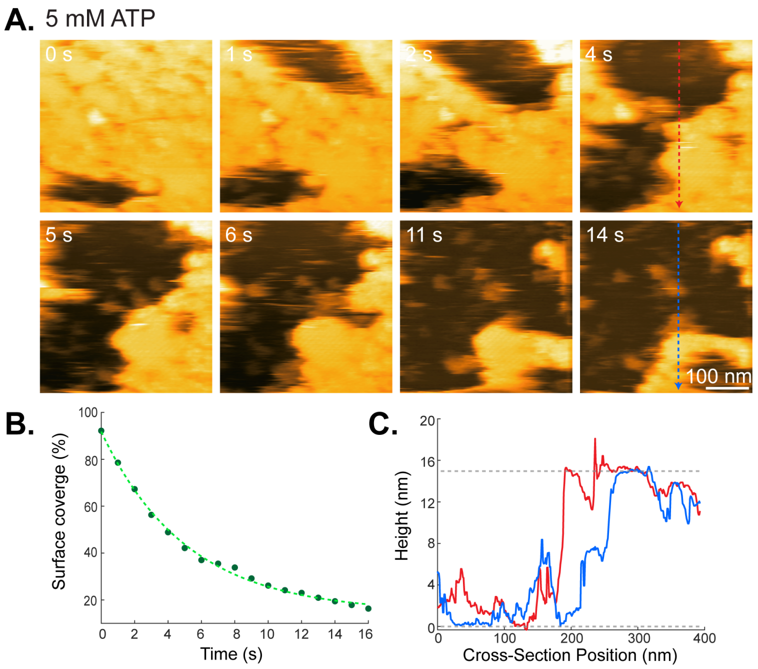

**Figure S3. Dissociation kinetics of ATP-treated PFKL WT lattice-like assemblies during HS-AFM imaging. (A)** Time-lapse HS-AFM images showing dissociation of the PFKL WT lattice-like assembly (225 nM protein in buffer containing 300 mM KCl and 5 mM ATP) during HS-AFM scanning. **(B)** Quantification of lattice dissociation based on surface coverage analysis, fitted with an exponential decay yielding a halftime of 3.5 s. **(C)** Representative cross-sectional height profiles across the mica substrate and the PFKL WT lattice-like assembly.

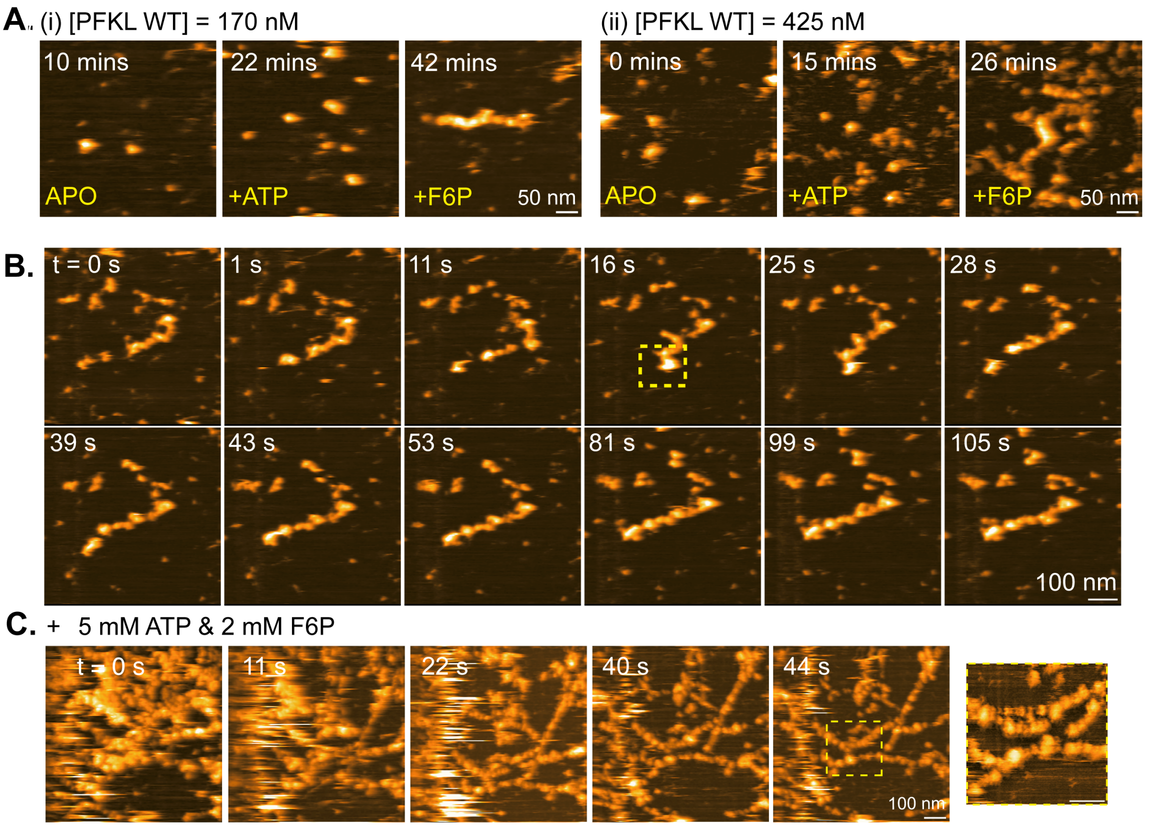

**Figure S4. PFKL WT self-assembles into filamentous structures upon F6P treatment. (A)** With a short incubation time on mica, PFKL WT formed small protein aggregates in both APO and ATP conditions. However, after adding 2 mM F6P into the 300mM KCl imaging buffer, PFKL WT self-assembled into filamentous structures, regardless of the (i) absence or presence of $Mg^{2+}$ ions. **(B)** Real-time HS-AFM imaging of PFKL WT (170 nM) showing filament formation after sequential addition of ATP and F6P in buffer containing 300 mM KCl. The resulting filaments remained stable during imaging without detectable dissociation. **(C)** Real-time observation of PFKL WT filament formation in the presence of 5 mM ATP and 2 mM F6P.

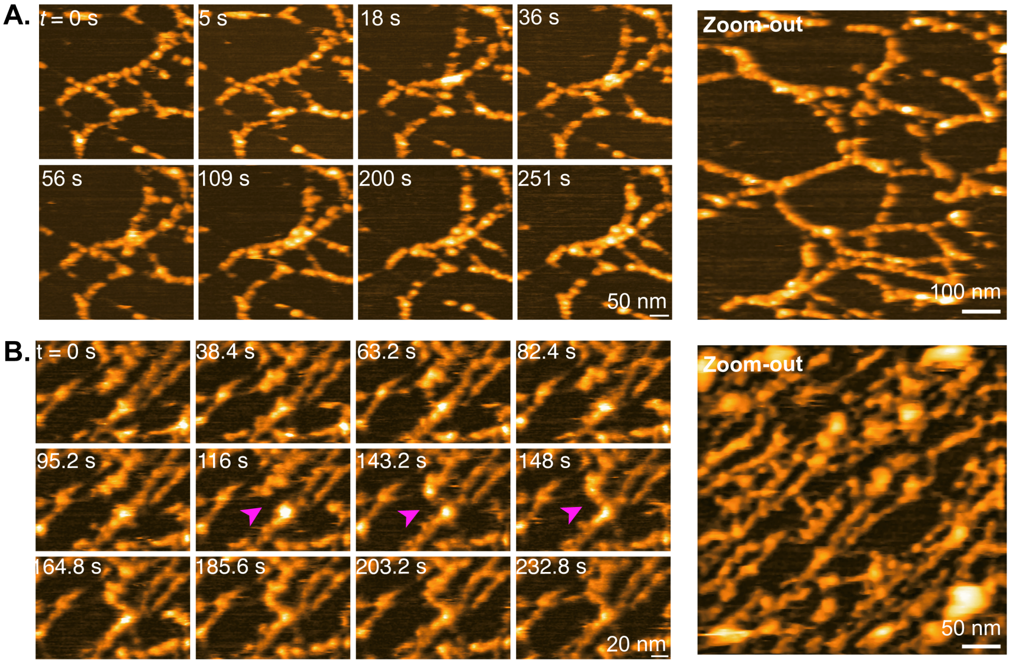

**Figure S5. PFKL WT filamentous structures in the presence of ATP and F6P. (A-B)** Real-time observations of PFKL WT filaments in network-like (protein concentration: 255 nM) and linear (protein concentration: 425 nM) geometry. During the observation, individual PFKL WT building blocks (tetramer or dimer of tetramers) exhibited dynamic movements and could locally form new filamentous connections (magenta arrows in **B**), but the overall filament architecture stayed unchanged**.**

242

243

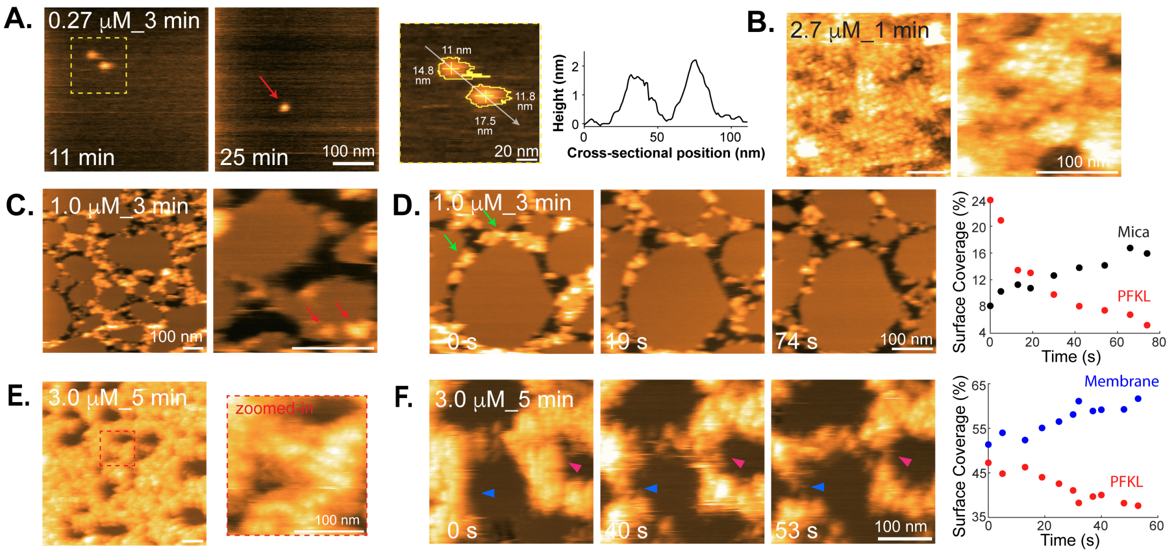

**Figure S6. PFKL binding and self-assembly on continuous membranes and membrane patches. (A-B)** Time-lapse HS-AFM images of APO-PFKL WT in 100 mM KCl buffer deposited onto a continuous, supported lipid bilayer (DOPC/DOPS=3/2). Insets indicate protein concentration and the pre-incubation time on the membrane (experimental conditions summarized in **Table S1**). At low protein concentration (0.27 μM), individual PFKL tetramers were observed bound to the membrane. At higher concentration (2.7 μM), APO-PFKL WT assembled into lattice-like assemblies on the membrane surface. **(C-F)** Time-lapse HS-AFM images of APO-PFKL WT in 100 mM KCl buffer deposited onto preformed membrane patches on mica. **(C, E)** Zoomed-out and zoomed-in views showing varied densities of PFKL WT localized on or adjacent to membrane patches. **(D, F)** Real-time observation of PFKL WT lattice dissociation at inter-patch gaps. Right panels: Quantitative analysis of surface coverage, distinguishing PFKL assemblies from mica or membrane background.

244

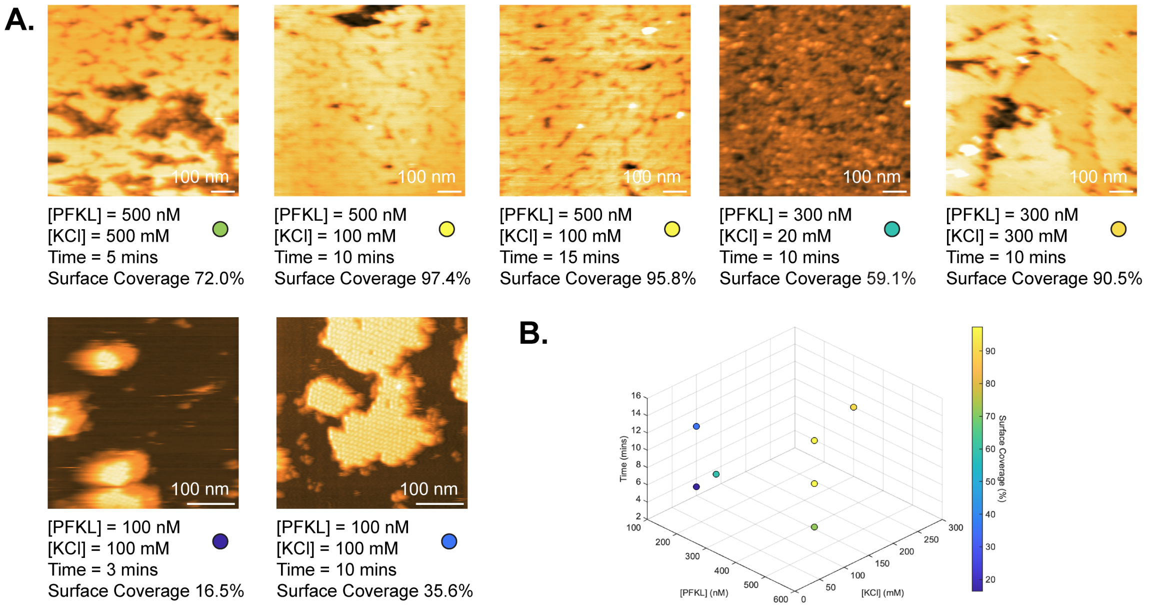

**Figure S7. PFKL N702T self-assembles into highly ordered double-layer 2D lattice under varied experimental conditions. (A)** Representative HS-AFM images of PFKL N702T 2D lattice formed on freshly cleaved mica at different protein and KCl concentrations and incubation times. When the KCl concentration was lowered to 20 mM, PFKL N702T self-assembled into relatively disorganized structures. **(B)** Surface coverage of PFKL N702T lattice as a function of protein concentration, incubation time, and KCl concentration, where the color represents the measured surface coverage percentage.

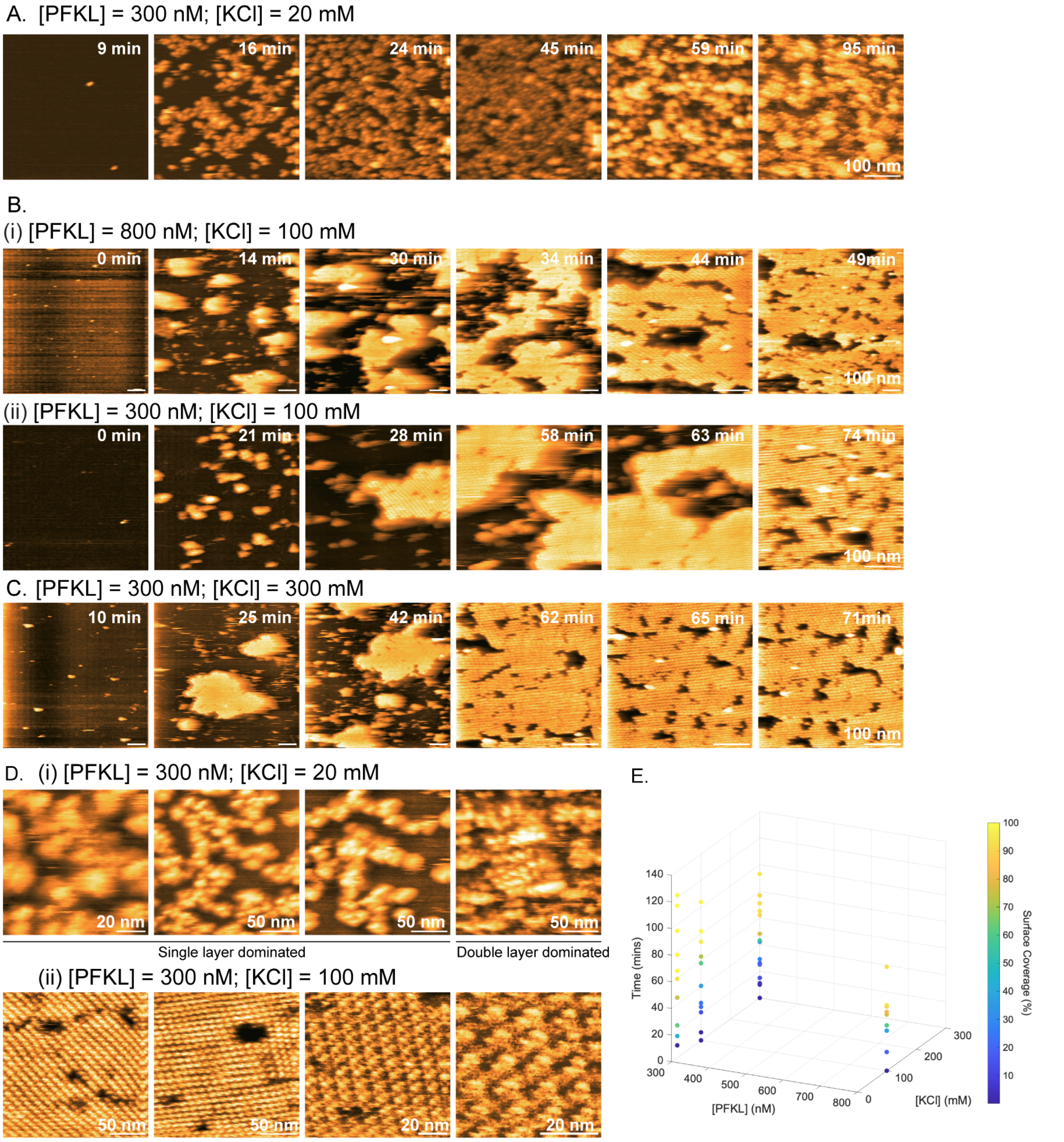

**Figure S8. Self-assembly of PFKL N702T under varied experimental conditions. (A-C)** Time-lapse traces of the PFKL N702T assembly process at different protein and KCl concentrations. **(A)** At 20 mM KCl, PFKL N702T self-assembled into a single layer and then formed a second layer with reduced long-range order. At higher ionic strength, **(B)** 100 mM and **(C)** 300 mM, PFKL N702T predominantly self-assembled as a double-layer lattice. The lattice islands expanded laterally and fused to form a continuous, double-layered lattice. **(D)** Zoomed-in high-resolution images of PFKL N702T self-assembly at (i) 20 mM KCl and (ii) 100 mM KCl concentration. **(E)** 3D plot of surface coverage of PFKL N702T assemblies at different protein concentration, KCl concentration, and incubation time.

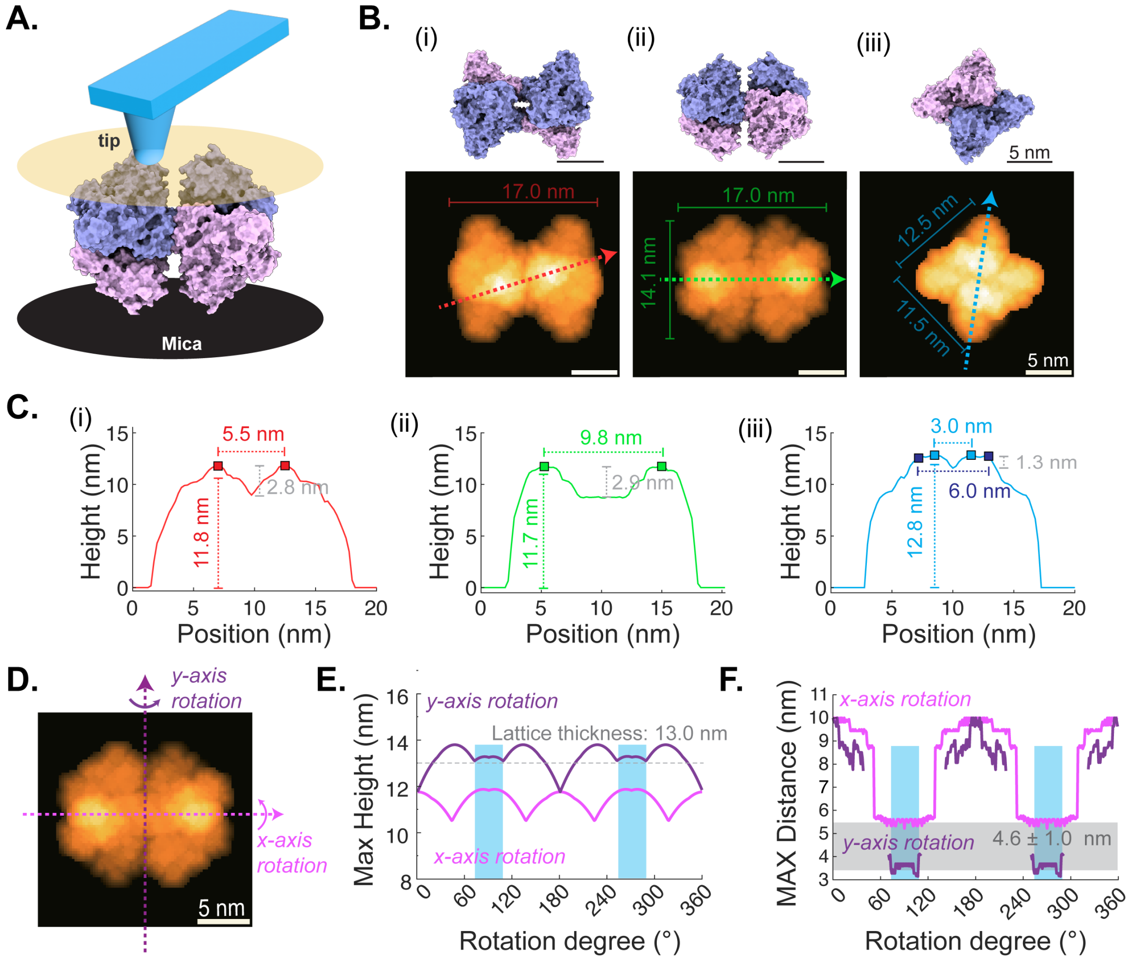

**Figure S9. AFM image simulation defines PFKL N702T tetramer orientation within the lattice. (A)** Simulated AFM imaging of the PFKL tetramer (PDB: 8W2G) using a computational probe with ~1.0 nm apex radius (15-degree tilt angle) to approximate HS-AFM tip convolution. **(B)** Simulated topographic projections of the tetramer from different orientations with labeled projected dimensions. **(C)** Representative cross-sectional profiles, extracted from (B), showing characteristic features, including protein height, distance between two protruding peaks, and peak-to-valley depth. **(D-F)** Systematic rotation of the tetramer along x- and y-axes and quantification of maximal height and peak separation. Experimental HS-AFM measurements (lattice thickness ~13.0 nm; peak spacing 4.6 ± 1.0 nm) match the simulated parameters of orientation (B-iii). In this configuration, the projected dimensions (~12.5 × 11.5 nm), thickness (~12.8 nm), peak spacing (~3-6 nm), and inter-peak depth (~1.3 nm) agree with lattice unit features observed in high-resolution HS-AFM images (**Fig. 5**).

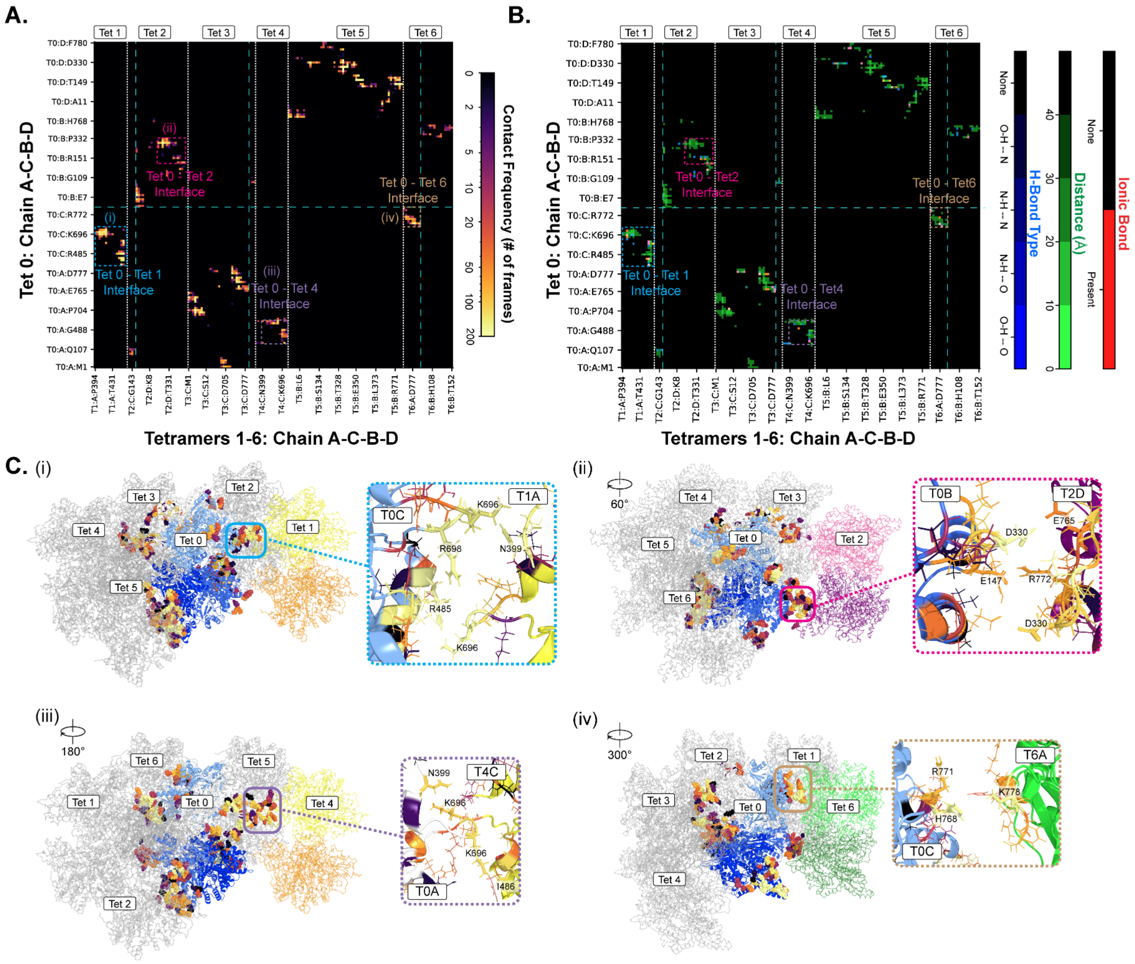

**Figure S10. Inter-tetramer contacts in the PFKL N702T lattice. (A-B)** Residue-residue contact maps showing **(A)** the frequency of inter-tetramer contacts observed during the 2-4 ns interval of the MD trajectory, with contacts defined by a Cβ-Cβ distance of 1.0 nm or less, and **(B)** a multichannel contact map of the inter-tetramer interaction network at 3 ns in the PFKL N702T lattice structure. The MD simulation trajectory is the same as that shown in **Fig. 6**. **(C)** Representative interfacial contacts at 3 ns in the PFKL N702T lattice structure, with residues making contacts colored according to contact frequency and shown as spheres. This panel shows additional representative inter-tetramer interfaces not highlighted in **Fig. 6E**.

252

### Description of Supplementary Movies

**Supplementary Movie 1. HS-AFM visualization of PFKL WT filament formation.** PFKL WT (270-425 nM) was introduced into the HS-AFM liquid chamber containing buffer with 300 mM KCl. In the presence of 1 mM ATP and 2 mM F6P, PFKL WT assembles into filamentous structures with distinct geometries, including networked and linear filaments (related to **Fig. 2G**). Scale bar: 100 nm. Imaging parameters: 1 frame/s, 2 nm/pixel (networked filaments); 1 frame/s, 1 nm/pixel (linear filaments).

**Supplementary Movie 2. APO-PFKL WT self-assembly on mica surfaces containing lipid membrane patches.** HS-AFM imaging of PFKL WT assembly on mica surfaces containing pre-deposited lipid membrane patches (DOPC/DOPS, 60:40) in buffer with 100 mM KCl (related to **Fig. 3B**). PFKL WT at the indicated concentrations was introduced into the HS-AFM liquid chamber. At lower protein concentrations, PFKL WT primarily assembled into lattices on exposed mica surfaces. At higher concentrations, lattice assemblies extended across and over the lipid membrane patches. Scale bar: 100 nm. Imaging parameters: 1 frame/s, 1 nm/pixel.

**Supplementary Movie 3. Edge dissociation of the PFKL N702T double-layer lattice.** HS-AFM imaging of a PFKL N702T (100 nM) lattice assembled on freshly cleaved mica after 10 min incubation in buffer containing 100 mM KCl (related to **Fig. 4A**). During imaging, edge dissociation exposes the underlying second lattice layer. Scale bar: 30 nm. Imaging parameters: 1 frame/s, 0.5 nm/pixel.

**Supplementary Movie 4. PFKL N702T self-assembles into a highly ordered 2D lattice at 100 mM KCl.** PFKL N702T (300 nM) was added to the HS-AFM liquid chamber in buffer containing 100 mM KCl (related to **Fig. 4D**, left). During HS-AFM imaging, PFKL N702T molecules freely diffuse and deposit onto freshly cleaved mica for self-assembly. Scale bar: 100 nm. Imaging parameters: 1 frame/s, 1 nm/pixel.

**Supplementary Movie 5. Self-assembly of PFKL N702T into an ordered 2D lattice at 300 mM KCl.** PFKL N702T (300 nM) was introduced into the HS-AFM liquid chamber containing buffer with 300 mM KCl (related to **Fig. 4D & S8C**). During imaging, individual PFKL N702T molecules diffuse in solution, adsorb onto freshly cleaved mica, and progressively assemble into a highly ordered two-dimensional lattice. Scale bar: 100 nm. Imaging parameters: 1 frame/s, 1 nm/pixel.

**Supplementary Movie 6. PFKL N702T assembly behavior at low ionic strength (20 mM KCl).** PFKL N702T (300 nM) was introduced into the HS-AFM liquid chamber containing buffer with 20 mM KCl (related to **Fig. 4E**). During imaging, individual PFKL N702T molecules diffuse to the mica surface and initially form a single-layer assembly. Subsequently, a secondary layer stacks on top of the first layer. The resulting structures exhibit a more island-like morphology, distinct from the well-ordered double-layer lattices observed at higher KCl concentrations. Scale bar: 100 nm. Imaging parameters: 1 frame/s, 1 nm/pixel.

**Supplementary Movie 7. High-resolution HS-AFM imaging of PFKL N702T lattices under different ligand conditions.** PFKL N702T (100–200 nM) was introduced into the HS-AFM liquid chamber containing buffer with 300 mM KCl (related to **Fig. 5A**). Lattice formation was examined under APO, 1 mM ATP, and 1 mM ATP + 2 mM F6P conditions. Across these conditions, no significant differences in higher-order lattice organization were observed between ligand-free and ligand-bound states. Scale bar: 100 nm. Imaging parameters: 1 frame/s, 1 nm/pixel.

**Supplementary Movie 8. High-resolution, zoomed-in HS-AFM imaging of PFKL N702T lattices under different ligand conditions.** PFKL N702T (100-200 nM) was introduced into the HS-AFM liquid chamber containing buffer with 300 mM KCl (related to **Fig. 5B**). Lattice structures were examined under APO, 1 mM ATP, and 1 mM ATP + 2 mM F6P conditions. At this higher magnification, individual lattice units are resolved, revealing intermolecular spacing and enabling analysis of tetramer orientation within the lattice. Scale bar: 10 nm. Imaging parameters: 0.6 frame/s, 0.33 nm/pixel (APO); 0.6 frame/s, 0.25 nm/pixel (ATP); 0.1 frame/s, 0.25 nm/pixel (ATP + F6P).

**Supplementary Movie 9. Progressive dissociation of PFKL N702T from the center of a double-layer lattice.** HS-AFM imaging of a highly ordered PFKL N702T two-dimensional lattice (100 nM) assembled on freshly cleaved mica after 10 min incubation in buffer containing 100 mM KCl (related to **Fig. 5E**). During imaging, individual PFKL N702T molecules gradually dissociate from the lattice center over time. Scale bar: 20 nm. Imaging parameters: 0.5 frame/s, 0.25 nm/pixel.

**Supplementary Movie 10. MD simulation of PFKL N702T lattice.** Representative video of MD simulation trajectory (2-4 ns) of the PFKL lattice, with front tetramers (Tet5 and Tet6) rendered transparent to highlight the packing geometry.
